# The saccadic repertoire of larval zebrafish reveals kinematically distinct saccades that are used in specific behavioural contexts

**DOI:** 10.1101/2023.11.07.565345

**Authors:** Charles K. Dowell, Joanna Y. N. Lau, Isaac H. Bianco

**Author notes:** Correspondence and Lead Contact: I.H.B.

## Abstract

Saccades are rapid eye movements that are used by all species with good vision. They have been extensively studied, especially in vertebrates, and are understood to be controlled by a conserved brainstem circuit. However, despite the fact that saccades play important roles during diverse visually guided behaviours, little is known about whether their properties, including the manner in which they are coordinated with head/body movements, vary in the context of different visuomotor tasks. Here, we characterise the saccadic repertoire of larval zebrafish and identify five saccade types, defined by systematic differences in kinematics and binocular coordination. Each type was differentially expressed during visually guided behaviours. Conjugate saccades form a large group that are used in at least four contexts: Fast phases of the optokinetic nystagmus, visual scanning in stationary animals, and to shift or maintain gaze during locomotion. Convergent saccades play a specialised role during hunting and are coordinated with body movements to foveate prey. Furthermore, conjugate and convergent saccades follow distinct velocity main sequence relationships and show differences in the millisecond coordination of the eyes and body, pointing to differences in underlying neurophysiology. In summary, this study reveals unexpected diversity in horizontal saccades and predicts saccade type-specific neural activity patterns.

**Highlights:**

- Kinematic analysis of thousands of rapid eye movements reveals five saccade types.
- Conjugate saccades have at least four identifiable visual functions.
- Convergent saccades are coordinated with body movements to foveate prey.
- Timing, kinematics and main sequence relationships indicate saccade type-specific neural control.

## Introduction

Saccades are brief but extremely rapid eye movements that are observed across species and phyla from crabs and cuttlefish to mice and primates (Land, 2019). They function to quickly shift the direction of gaze between stable fixations and intermittently recentre the eyes during compensatory nystagmus (vestibuloocular and optokinetic reflexes that operate to minimise retinal image slip). Most species coordinate saccades with head rotations. However, they are also used independently of head movements in foveate animals to successively shift the point of fixation for high-spatial frequency sampling of the visual environment (Yarbus, 1967; Robinson, 2022b). Much is known about the kinematic properties of saccades and their underlying neurophysiology (Sparks, 2002; Robinson, 2022c). Although properties such as latency, duration and velocity can vary as a function of whether saccades are made when the head is free to move versus restrained (Meyer et al., 2020), in the dark versus light (Sharpe et al., 1975), or directed to visible versus remembered targets (Smit et al., 1987), it is generally considered that in vertebrates all saccades are generated by a common, evolutionarily conserved, brainstem circuit. However, saccades play a role in a wide array of visually guided behaviours and little is known about whether there might be systematic differences in their kinematics and/or patterns of coordination with other body movements across different behavioural contexts, perhaps even to the extent that different subtypes of saccade can be recognised. This represents an important knowledge gap, because understanding a species’ motor repertoire is essential for identifying sensorimotor rules that underly more complex behaviour, in turn providing important insights into underlying neural computations.

The larval zebrafish is an important model in neuroscience research (Friedrich et al., 2010) and has been used to study the development and neural control of oculomotor behaviours. Most studies of saccades have focussed on spontaneous conjugate saccades and fast phases of the optokinetic reflex (OKR) in restrained animals and have revealed neural activity that controls the timing and direction of spontaneous saccades (Ramirez & Aksay, 2021; Wolf et al., 2017), optogenetically mapped the locus of saccade generation in rhombomere 5 (Schoonheim et al., 2010), and described monocular and binocular encoding in saccade-active cells (Leyden et al., 2021). In addition, studies of prey-catching behaviour have shown that zebrafish initiate hunting routines using a convergent saccade and a high ocular vergence angle is then sustained throughout prey-tracking (Bianco et al., 2011; Trivedi & Bollmann, 2013). However, the extent to which these encompass the diversity of saccadic eye movements is unknown and there has been little examination of rapid eye movement kinematics.

Here we set out to describe the full repertoire of saccades of larval zebrafish by measuring rapid eye movements in tethered and freely swimming animals engaged in a range of visuomotor behaviours. We distinguished five major saccade types, defined by systematic differences in kinematics and binocular coordination and found they were differentially engaged across different behavioural contexts. Conjugate saccades formed a large group with four identifiable visual roles, including two distinct patterns of coordination with head/body rotations whereas convergent saccades mediated goal-directed orientations to foveate prey targets during hunting. High temporal-resolution recordings revealed that conjugate and convergent saccades differed in the timing of binocular eye movements and eye-body coordination and remarkably, conformed to different velocity main sequence relationships, indicating they are controlled by distinct patterns of neural activity. Overall this study provides insight into the visuomotor behavioural strategies used by larval zebrafish and motivates hypotheses about circuit control of saccadic eye movements and eye-body coordination.

## Results

### Characterisation of the saccadic repertoire of larval zebrafish

We first sought to estimate the full repertoire of saccadic eye movements of larval zebrafish by tracking the behaviour of both freely swimming and tethered fish engaged in a range of behaviours. Tethered larvae were restrained using low-melting point agarose, but with sections cut away to permit free movement of the eyes and tail [Figure 1A]. They were presented with a range of visual stimuli that included whole-field drifting gratings, which evoke optokinetic nystagmus (OKR, comprising slow phase rotations in the direction of visual motion with intermittent fast ‘reset’ saccades) (Huang & Neuhauss, 2008) and optomotor swimming (Neuhauss et al., 1999; Orger et al., 2000), small, prey-like moving spots which evoke hunting responses involving saccadic eye convergence (Bianco et al., 2011), as well as dark-flashes and looming stimuli that evoke high-angle turns and avoidance swims (Burgess & Granato, 2007; Dunn et al., 2016a).

**Figure 1:**
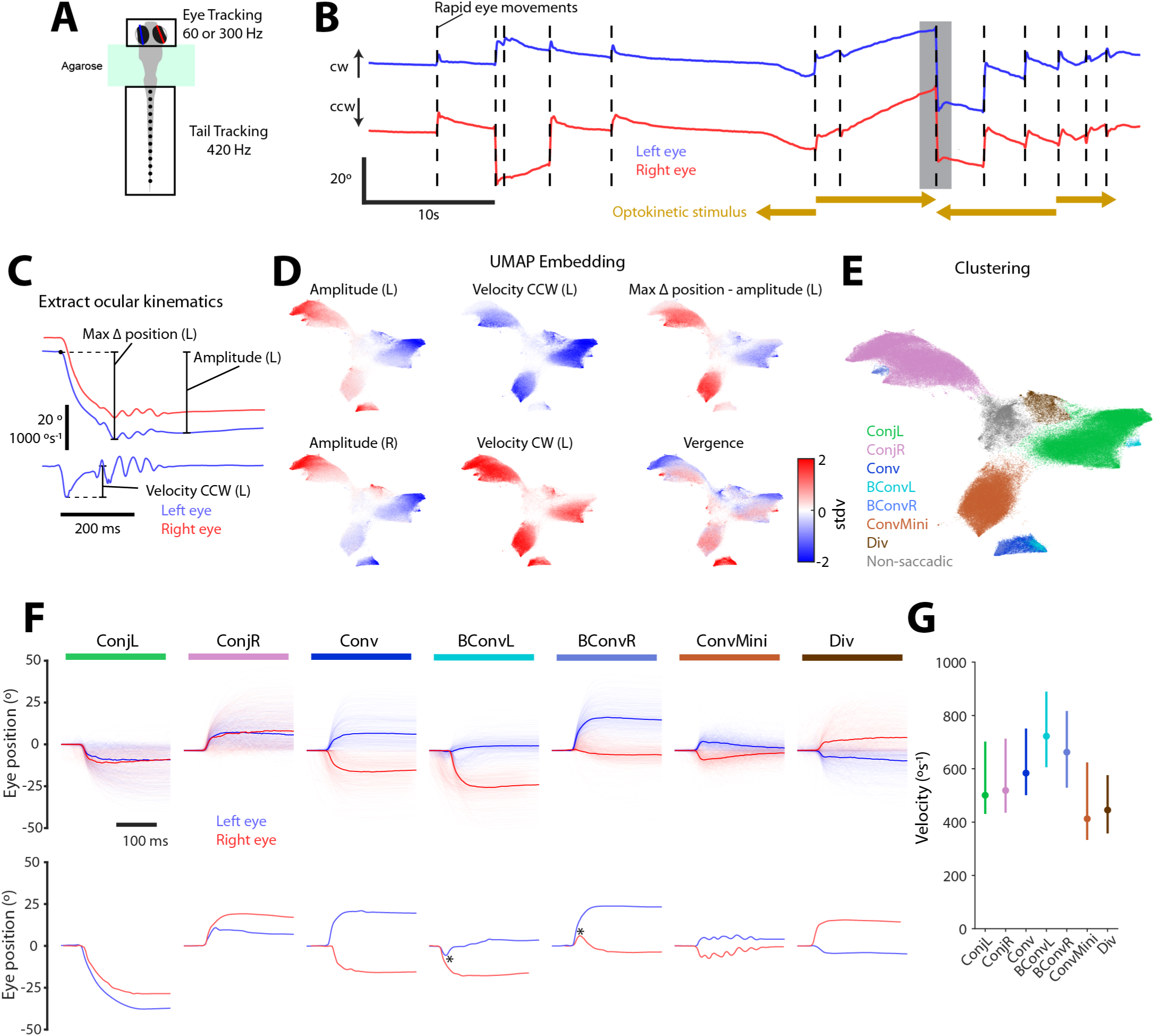
Saccade detection and classification. (A) Illustration of tethered behavioural tracking. (B) Eye position time series data from an example 60 s experimental epoch. Upwards corresponds to clockwise (left eye nasal, right eye temporal) rotation. Automatically detected rapid eye movement events indicated with dashed lines. In this epoch, leftwards and rightwards drifting gratings were presented in front of the larva to evoke optokinetic nystagmus, as indicated. (C) A subset of position (top) and velocity (bottom) metrics for an example rapid eye movement (time of this event indicated by grey shading in B). Letter in parentheses indicates left or right eye. See Methods for details. (D) Rapid eye movements (213,462 events from 152 animals) embedded in 2D UMAP space and coloured by normalised oculomotor metrics. Additional metrics shown in Figure S1. (E) Rapid eye movements embedded in 2D UMAP space and coloured by saccade type label. (F) *Top:* For each saccade type, 500 eye position traces are plotted with the median overlaid in bold. *Bottom:* Single example saccades. * indicates reversal of eye velocity during biphasic convergent saccades. (G) Eye velocity for all saccade types. Data plotted as median (±IQR) across mean values from each larva (*N* = 152). *See also Figure S1 and Figure S2*.

We described the kinematic features of rapid eye movements, initially focussing on datasets from tethered animals where we could obtain very high-quality tracking data (*N* = 152 larvae, 6–7 days post fertilisation). Putative rapid eye movements were first detected as peaks in the convolution of the eye-in-head position (hereafter eye position) time series and a step filter [Figure 1B]. For each event, we then computed nine position and velocity metrics [Figure 1C; Methods] describing kinematic features of both the left and right eye. We note that putative saccadic events were characterised in terms of movement of both eyes to enable assessment of different patterns of binocular coordination. Kinematics included the amplitude and peak velocity of each eye’s movement, post-saccadic vergence and a metric describing the extent to which each eye’s position was maintained after the putative saccade (the difference between max delta position and amplitude). Next, we used UMAP to embed *N* = 213, 462 events in two dimensions and observed a smooth and systematic variation of kinematic values across the embedding space [Figure 1D; Figure S1A]. Several regions appeared to be quite well separated from one another and contained rapid eye movements with similar distributions of kinematic features. We therefore applied a density based clustering procedure to the embedded data and thereby classified rapid eye movements using seven cluster labels [Figure 1E]. Clusters had distinct and unimodal kinematic distributions [Figure S1B-D], indicating this classification scheme captured the major patterns of variation across the dataset.

The seven labels defined five saccade types (two types were subdivided into left- and right-directed clusters), which included both conjugate and disjunctive eye rotations, wherein left and right eyes moved either in the same or opposite directions, respectively [Figure 1F]. Conjugate saccades were assigned to two large clusters (left- and right-directed ‘Conj’), within which there was continuous variation in kinematic properties [Figure 1D, Figure S1]. Convergent saccades, in which both eyes rotate nasally, fell into four clusters: Regular convergent saccades (‘Conv’) and biphasic convergent saccades (left- and right-directed ‘BConv’) both resulted in large and sustained elevations in vergence [Figure S1D]; biphasic saccades had the distinctive feature that one eye first made a small temporal rotation before reversing direction and rotating nasally [Figure 1F, bottom]. Zebrafish also generated a large number of miniature convergent saccades (‘ConvMini’) involving a small, transient increase in vergence that decayed rapidly. Finally, we observed a small number of divergent saccades (‘Div’). Eye velocity varied systematically across saccade types, with median values ranging from 400 to over 700*^◦^/s* [Figure 1G].

We could identify the same saccade types in freely swimming larval zebrafish. To show this, we processed eye tracking data to detect rapid eye movements, extracted the same nine kinematic metrics, and then performed a supervised low-dimensional embedding using the tethered UMAP solution as a template (*N* = 9, 367 events from 8 animals). Despite the fact that this was a smaller dataset, rapid eye movements from freely swimming animals spanned the kinematic embedding space and could be classified with the same seven labels [Figure S2A]. Inspection of eye position traces confirmed these saccades showed similar features, including patterns of binocular coordination, as compared to the saccades of tethered animals [Figure S2B].

### Saccade types are used in distinct contexts

Saccadic eye movements serve a variety of visual functions and are often coordinated with head and body movements to redirect gaze. Therefore, we next investigated how the different saccade types of larval zebrafish are utilised in different behavioural contexts.

Conjugate saccades comprised the most frequent type of rapid eye movement (*∼* 70% of all saccades in both tethered and freely swimming animals, [Figure 2A]). As was the case for all saccade types, they were usually accompanied by a swim bout [Figure 2B], but were also produced by stationary animals [Figure 2C-D]. When either tethered or freely swimming animals generated left or right turns, we observed an elevated frequency of conjugate saccades of the same laterality, suggesting these rapid eye movements contribute to combined eye-body gaze shifts, in line with previous observations (Wolf et al., 2017). When presented with drifting gratings designed to evoke the optokinetic response, larvae generated conjugate saccades, unaccompanied by swims, in the opposite direction to whole-field motion, indicating these are fast phases of the optokinetic nystagmus, serving to recentre eye position in the orbit [Figure 2C].

**Figure 2:**
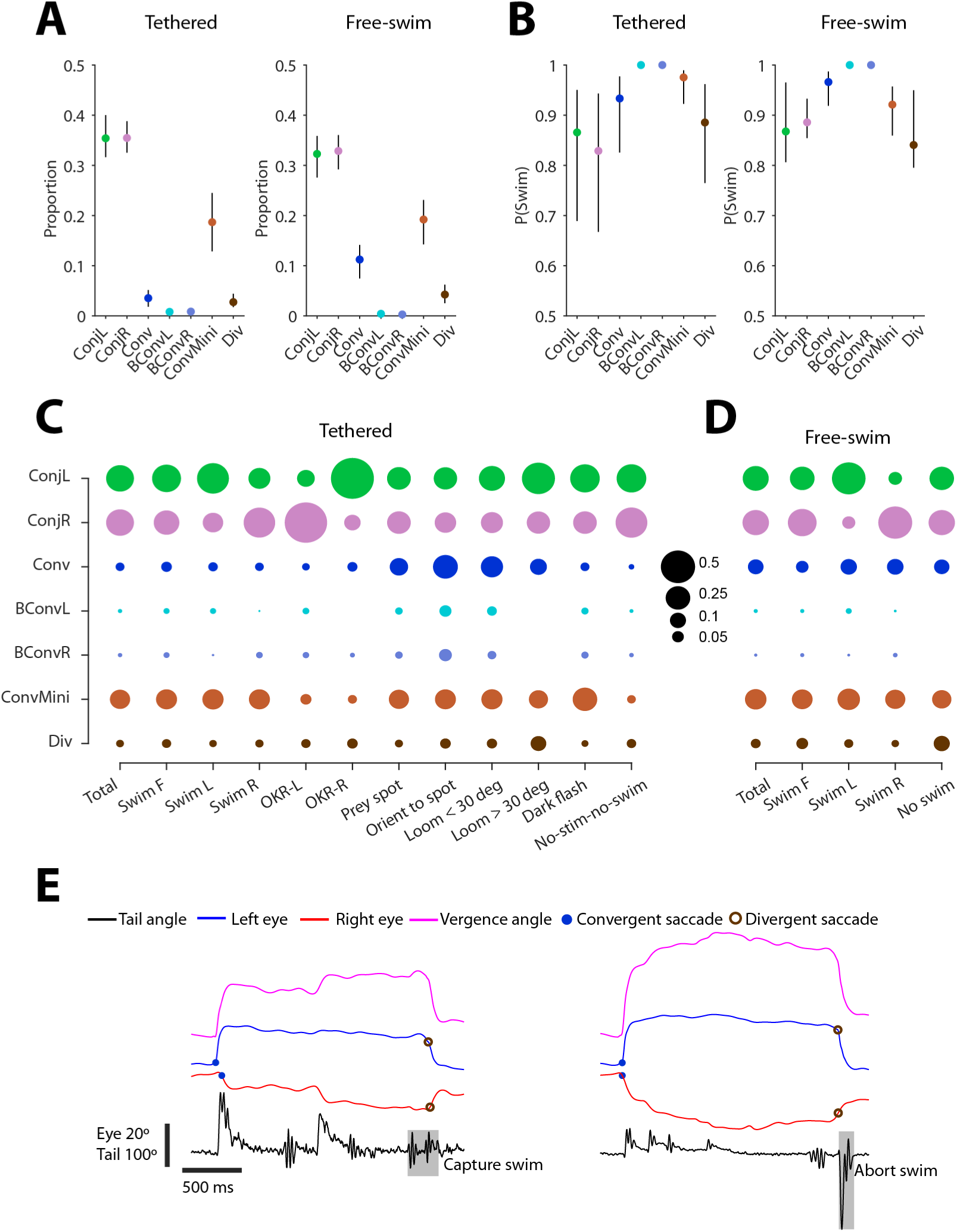
Contextual deployment of saccades. (A) Proportion of each saccade type for tethered and free-swimming datasets (median ± IQR across *N* = 152 tethered and *N* = 8 free-swimming larvae). (B) Probability of a swim occurring within 200 ms of a saccade, for each saccade type. (C) Proportion of each saccade type observed during the indicated contexts for tethered larvae. Swim F, forward swim; Swim R/L, right/left swim (abs. tail bend angle ≥25◦); OKR-L/R, left/rightwards OKR grating and no swim; Prey spot, small prey-like moving spot; Orient to spot, first orienting turn to prey-like stimulus. (D) Proportion of each saccade type during the indicated contexts for free-swimming larvae. Swim F, forward swim; Swim R/L, right/left swim (abs. orientation change ≥10◦). (E) Examples of hunting sequences that end with a divergent saccade coincident with either a capture swim (left) or an abort swim (right).

Convergent saccades are a defining feature of hunting behaviour (Bianco et al., 2011; Trivedi & Bollmann, 2013) and in accordance with previous studies, we observed an elevated frequency of both regular and biphasic convergent saccades when larvae were presented with, and oriented towards, small prey-like moving spots [Figure 2C]. Convergent saccades also occurred ‘spontaneously’, at a low rate (Bianco & Engert, 2015; Zylbertal & Bianco, 2023), and were quite frequently evoked by looming stimuli, especially early in stimulus presentation while the expanding spot was small (*<* 30*^◦^*) and presumably perceived as a prey-like stimulus.

Miniature convergent saccades were observed in most contexts [Figure 2C,D], making their role rather unclear. However, they were rarely observed in tethered, stationary larvae. Divergent saccades were uncommon, but occurred at elevated frequency in response to looming stimuli (*>* 30*^◦^*), compatible with a role in redirecting gaze behind the animal in combination with the high-angle avoidance turns elicited by this stimulus (Dunn et al., 2016a; Marques et al., 2018). They also occur at the end of hunting sequences to switch out of the high-vergence predatory mode of gaze; thus we observed divergent saccades after larvae performed capture strikes or aborted prey-tracking [Figure 2E].

Because conjugate (Conj) and convergent (Conv, BConv) saccade types spanned a similar range of amplitudes but showed substantial variation in their utilisation in different behavioural contexts, we focussed on these types for the remainder of the study. In particular, we characterised how these saccades are coordinated with swims to generate gaze shifts and compared their detailed kinematics to make inferences about underlying circuit activity.

### Conjugate saccades show two patterns of coordination with swims to enable gaze-shifting and gaze-maintenance

Across many species, planned gaze shifts are accomplished by coordinated eye and head movements, but in some species saccades alone can redirect gaze when the head/body is stationary (Robinson, 2022b). Because conjugate saccades in zebrafish occurred both with and without accompanying swims, we next explored the ways in which they contribute to gaze changes.

Zebrafish larvae generated their largest amplitude conjugate saccades when they were stationary. This was evidenced by plots of saccade amplitude coded by the probability of a coincident swim (defined as swim bouts within 200 ms of saccade onset), which showed that, in tethered animals, conjugate saccades exceeding 20*^◦^* were rarely accompanied by body movement [Figure 3A]. This relationship was also observed under freely swimming conditions [Figure S3A], albeit at lower frequency because larvae that were free to move spent little time stationary. These observations indicate that zebrafish can redirect gaze using saccadic eye movements alone and that when they are stationary, large amplitude conjugate saccades allow them to maintain visual exploration and scan their environment.

**Figure 3:**
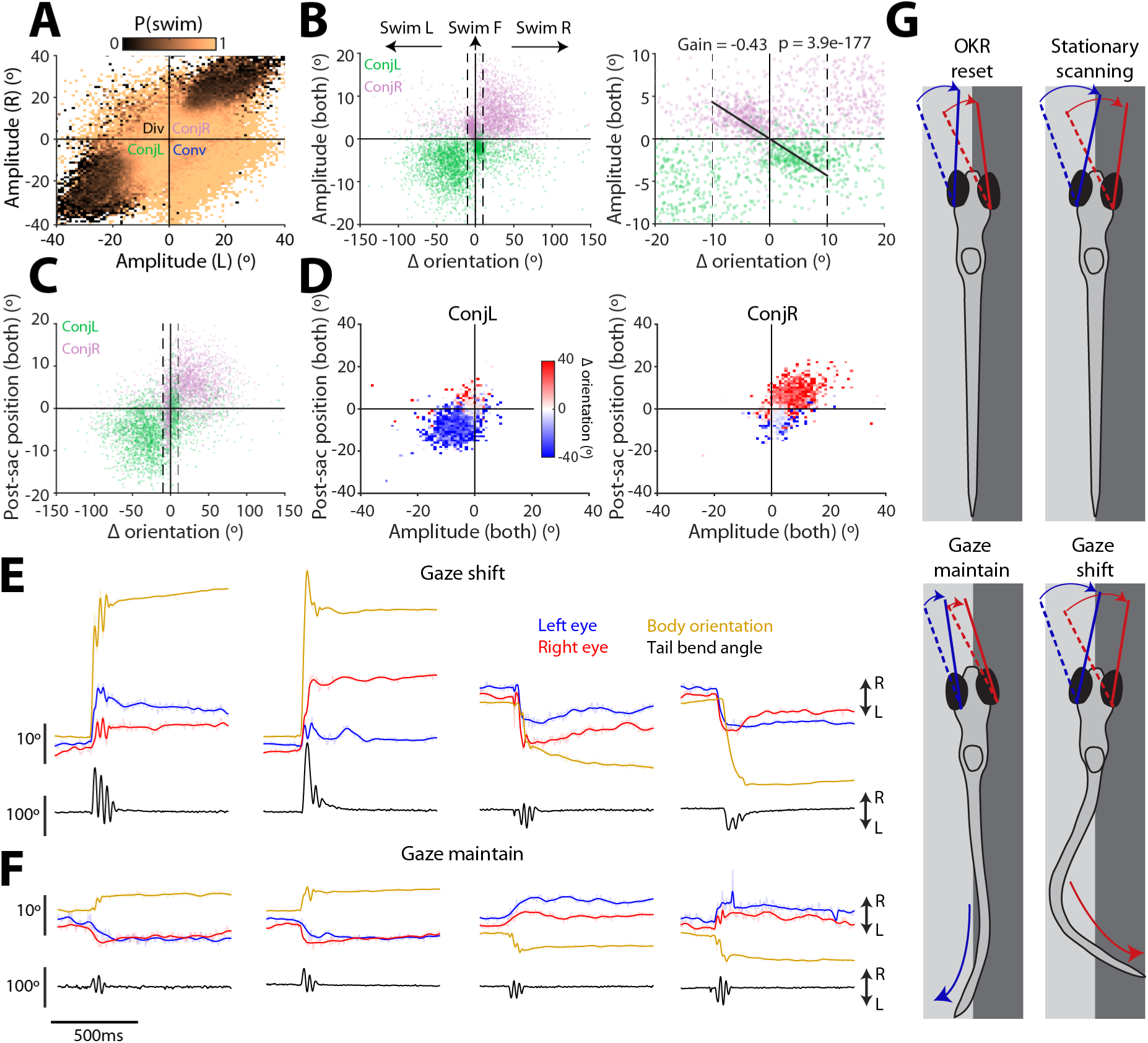
Conjugate saccades are produced by stationary animals and are coordinated with swims to either shift or maintain gaze. (A) Saccade amplitude coded by the probability of a coincident swim (data from 152 tethered larvae). (B) *Left:* Change in conjugate eye position (mean of left- and right-eye amplitude) versus change in body orientation (5,869 saccades from 8 free-swimming fish). Dashed lines indicate thresholds for forward swims (–10 ≤ Δori ≤ 10*^◦^*) versus turns. *Right:* Magnified portion of left panel highlighting gaze-maintaining conjugate saccades. Linear fit calculated for saccades with direction opposite to body reorientation and where |Δ*ori*| *<* 10*^◦^*(gradient = 0.43, *R*^2^ = 0.39, *N* = 820 saccades, p-value t-test). (C) Post-saccadic eye position (mean of left- and right-eye position) versus change in body orientation. (D) Left and right conjugate saccades binned by amplitude and post-saccadic eye position and colour-coded by median change in body orientation. (E–F) Examples of gaze-shifting (E) and gaze-maintaining (F) conjugate saccades. Smoothed eye position is plotted in bold over raw data. (G) Summary of four contexts in which larval zebrafish use conjugate saccades. *See also Figure S3*.

Next, we examined the kinematics of saccades that were coincident with swims and observed two distinct relationships between eye and body reorientations. As described above, left- and right-directed turns were frequently accompanied by conjugate saccades that shifted gaze in the same direction [Figure 2]. By plotting the change in body orientation of freely swimming larvae versus the amplitude of the conjugate eye movement (the mean of left- and right-eye amplitude), we observed that for body orientation changes exceeding 10*^◦^*, the majority (83.0 *±* 3.6%*, N* = 5, 869) were accompanied by conjugate saccades of the same laterality [Figure 3B]. Although conjugate amplitude was weakly correlated with body reorientation (Pearson’s rho = 0.09, *R*^2^ = 0.056, *p* = 2.1*e −* 09 t-test) [Figure 3B, upper right and lower left quadrants] and the overall gaze shift (Pearson’s rho = 0.10, *R*^2^ = 0.040, *p* = 1.1*e −* 10) [Figure S3C], eye position following the gaze shift was consistently displaced in the direction of locomotion [Figure 3C]. Specifically, post-saccadic eye position matched turn direction for 85.4 *±* 4.0% of gaze shifts (for which absolute change in body orientation exceeded 10*^◦^*). We also showed this by colour-coding post-saccadic eye position by body reorientation [Figure 3D] (or swim lateralisation for tethered animals [Figure S3B]) and found that rightwards conjugate saccades that terminated rightwards of primary eye position were associated with rightwards swims. To summarise, zebrafish shift their gaze using combined eye-body movements and orient their visual field in the direction in which they are moving [Figure 3E].

Zebrafish also displayed a second pattern of eye-body coordination, in which conjugate saccades were paired with swims of the *opposite* laterality [Figure 3B,C]. For instance, rightwards conjugate saccades were sometimes coincident with leftwards body movements. This was especially evident when changes in body orientation were small, *<* 10*^◦^*, within a range that can be considered as ‘forward swims’ (Naumann et al., 2016) [Figure 3B right]. In these instances, the magnitude of the change in conjugate eye position was approximately half the change in body orientation resulting from the swim (*gain* = *−*0.43, *R*^2^ = 0.39, Figure 3B right). That these eye movements were saccades, as opposed to oscillations produced by compensatory spino-ocular coupling (Straka et al., 2022), was evidenced by both raw eye tracking data [Figure 3F] as well as analysis of velocity main sequence relationships (see below). Thus, zebrafish use small amplitude conjugate saccades to compensate for body rotation and thereby stabilise vision during forward locomotion.

In summary, larval zebrafish use conjugate saccades in four behavioural contexts [Figure 3G]: 1) Fast phases of the optokinetic nystagmus serve to recentre eye position; 2) Large amplitude saccades are generated by stationary animals, likely subserving visual exploration; 3) Saccades coincident with body turns of the same laterality are used to shift gaze; 4) Small saccades with laterality opposite to body rotation help to maintain gaze direction during forward locomotion.

### Convergent saccades enable precise gaze shifts that foveate prey

A defining feature of zebrafish hunting behaviour is that all hunting routines commence with a convergent saccade and a high vergence angle is then sustained throughout prey-tracking (Bianco et al., 2011; Trivedi & Bollmann, 2013). It has been suggested that by increasing the extent and proximity of the binocular visual field this may support a simple stereopsis mechanism for judging distance to prey at the moment immediately prior to capture strikes (Bianco et al., 2011). However, when larvae first initiate hunting, convergent saccades are often lateralised towards prey (Trivedi & Bollmann, 2013; Henriques et al., 2019), raising the possibility that they help to binocularly visualise the target from the first orienting response. We therefore examined how saccades are coordinated with body movements in the context of goal-directed orientations to prey.

Zebrafish initiated hunting routines using both regular and biphasic convergent saccades [Figure 4A]. Regular convergent saccades were used when prey was located in the anterior visual field (mean azimuth = 0.4*^◦^*, mean absolute azimuth = 52.9*^◦^*), whereas larvae responded to more eccentric prey with left- or right-directed biphasic convergent saccades (mean azimuth = 1.8*^◦^*, mean absolute azimuth = 88.2*^◦^*). Convergent saccades more than doubled the size of the binocular visual field (from 19 to 47*^◦^*) [Figure 4B] and post-saccadic conjugate eye position smoothly covaried with body reorientation, such that larger turns were associated with more lateralised egocentric gaze [Figure 4C]. Thus, convergent saccades have both vergence and version components that increase the extent of the binocular visual field and horizontally shift gaze in cooperation with body movements, respectively.

**Figure 4:**
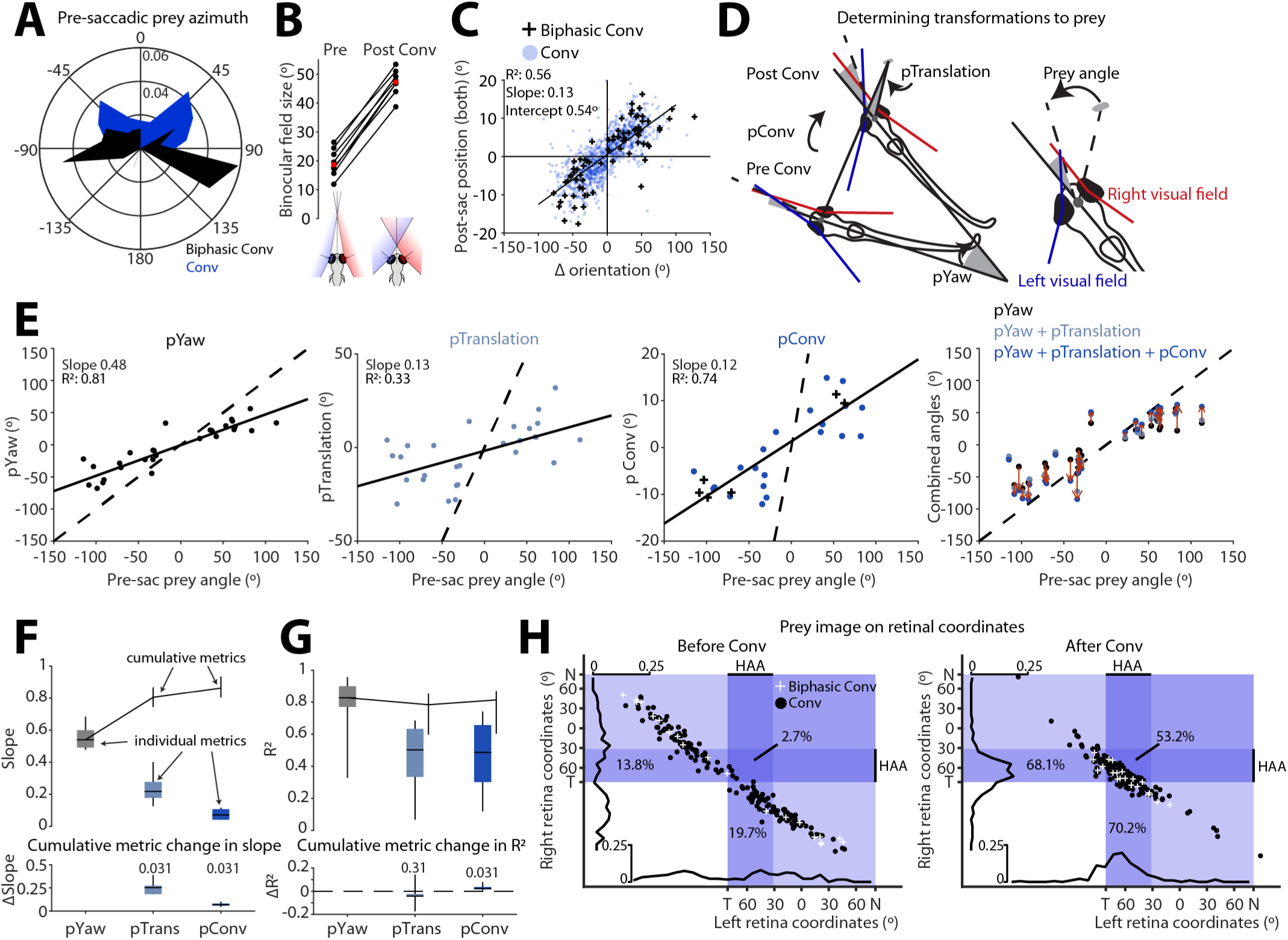
Convergent saccades foveate prey from the onset of hunting. (A) Prey azimuth at the time of the convergent saccade that defines the onset of a hunting epoch (202 Conv and 25 BConv from 8 fish). (B) Convergent saccades increase the size of the binocular visual field. Median binocular field size pre- and post-saccades for *N* = 8 fish. Median across animals in red. (C) Post-saccadic conjugate eye position (mean of left- and right-eye position) versus change in body orientation, with least squares regression fit (*N* = 1178 saccades from 8 fish). (D) Schematic illustrating how pYaw, pTrans and pConv are computed for the orienting response to prey. Angles corresponding to each metric shown by grey shaded regions. *Right:* Schematic illustrating gaze-referenced prey position, defined as the angle between the vectors connecting the midpoint between the eyes to (i) the prey target and (ii) the nearest point of binocular overlap. (E) pYaw, pTrans and pConv versus pre-saccadic prey position for the first orienting responses of 28 hunting epochs from one example fish. Linear fits, with slope and *R*^2^, shown as solid lines. Rightmost panel shows combined eye-body orienting response (by iteratively summing pYaw, pTrans and finally pConv); red arrows indicate change from pYaw to full response. In all panels *y* = *x* shown as dashed line. (F-G) *Upper panels:* Slope (F) and goodness-of-fit (G) of regression fits to individual (box-plots) and cumulative (shown as median ± IQR) prey orientation metrics versus pre-saccadic prey angle. *Lower panels:* Change in slope (F) and goodness-of-fit (G) with addition of pTrans followed by pConv. p-values signed rank test. *N* = 6 fish. (H) Prey image in naso-temporal retinal coordinates before (left) and after (right) the first orienting response (188 hunting sequences from 6 fish). Each eye is assumed to have a field of view of 163*^◦^* (Easter Jr & Nicola, 1996) and a high acuity area spanning 50*^◦^* of temporal retina (Yoshimatsu et al., 2020). Percentage of prey targets seen by HAA of one or both eyes are shown.

To assess how larvae control convergent saccades and accompanying body movements in the context of goal-directed predatory gaze shifts, we decomposed the first orienting response towards prey (188 individual hunting epochs from 6 animals). We mapped prey position (angle in the horizontal plane) to a gaze-referenced coordinate system [Figure 4D, inset] and evaluated the specific contributions of body rotation, body translation and the saccadic eye movement to redirecting gaze towards the target [Figure 4D]. We found that zebrafish smoothly modulated all three components in accordance with prey position [Figure 4E-F]. Body rotation (pYaw) made the largest contribution to the change in gaze-referenced prey position, with a magnitude of approximately half of initial prey azimuth (*gain* = 0.56 *±* 0.03*, N* = 6 fish). Translation of the head resulting from the first swim bout (pTrans) further served to align the frontal axis with the prey target, with gain of 0.24 *±* 0.04. Finally, the saccadic eye movement provided an additional goal-directed gaze shift (pConv, *gain* = 0.08 *±* 0.01). As a result, the combined eye-body movement shifted binocular gaze towards the target prey with an overall gain of 0.88*±*0.04.

To estimate how these three behavioural components impact the precision of the orienting response, we assessed the goodness-of-fit (*R*^2^) of linear fits to prey position, while successively including the pYaw, pTrans and pConv components of individual reorientations [Figure 4G]. As well as having the highest gain, pYaw was the most precise motor component, having a mean *R*^2^ of 0.87 *±* 0.03. The pTrans component was less accurate (*R*^2^ = 0.50 *±* 0.06), indicating greater stochasticity in body displacement. Interestingly, the addition of pConv resulted in a small but significant increase in orientation accuracy (Δ*R*^2^ = 0.01 *±* 0.008*, p* = 0.031). Thus, convergent saccades act in cooperation with body movements during goal-directed visual orientations towards prey and may compensate for errors attributable to swimming movements.

Next, we tested the hypothesis that the high gain of the eye-body orienting response is sufficient to binocularly foveate prey from the onset of hunting. Zebrafish larvae have a fovea-like high acuity area (HAA) in the ventral-temporal retina with an elevated density of UV cones that is thought to be crucial for visualising UV-scattering prey (Schmitt & Dowling, 1999; Yoshimatsu et al., 2020). By projecting prey location into retinal coordinates [Figure 4H], we estimated that for the vast majority of hunting epochs the first orientating movement shifted the image of prey to the HAA of at least one eye (84.3 *±* 3.1%, *N* = 6 fish); moreover, in half the epochs, this first eye-body manoeuvre was sufficient to binocularly foveate prey (50.3*±*4.7%).

In sum, convergent saccades are coordinated with body movements to allow zebrafish to accurately foveate their prey from the onset of hunting.

### Distinct timing rules for binocular and eye-body coordination across saccade types

When coordinated eye and body movements are used to redirect gaze, saccades typically precede head/body rotations by a few milliseconds. However, relative movement timing depends on several factors, including task requirements (Freedman, 2008). Because conjugate and convergent saccades were coordinated with swims to redirect gaze in distinct behavioural contexts, we examined movement timing to gain insight into whether there might be differences in the organisation and coordination of the underlying neural commands. For this analysis, we used high temporal resolution (300 Hz) tracking data from 58 tethered larvae to estimate inter-ocular and eye-tail latencies ([Figure 5A], Methods).

**Figure 5:**
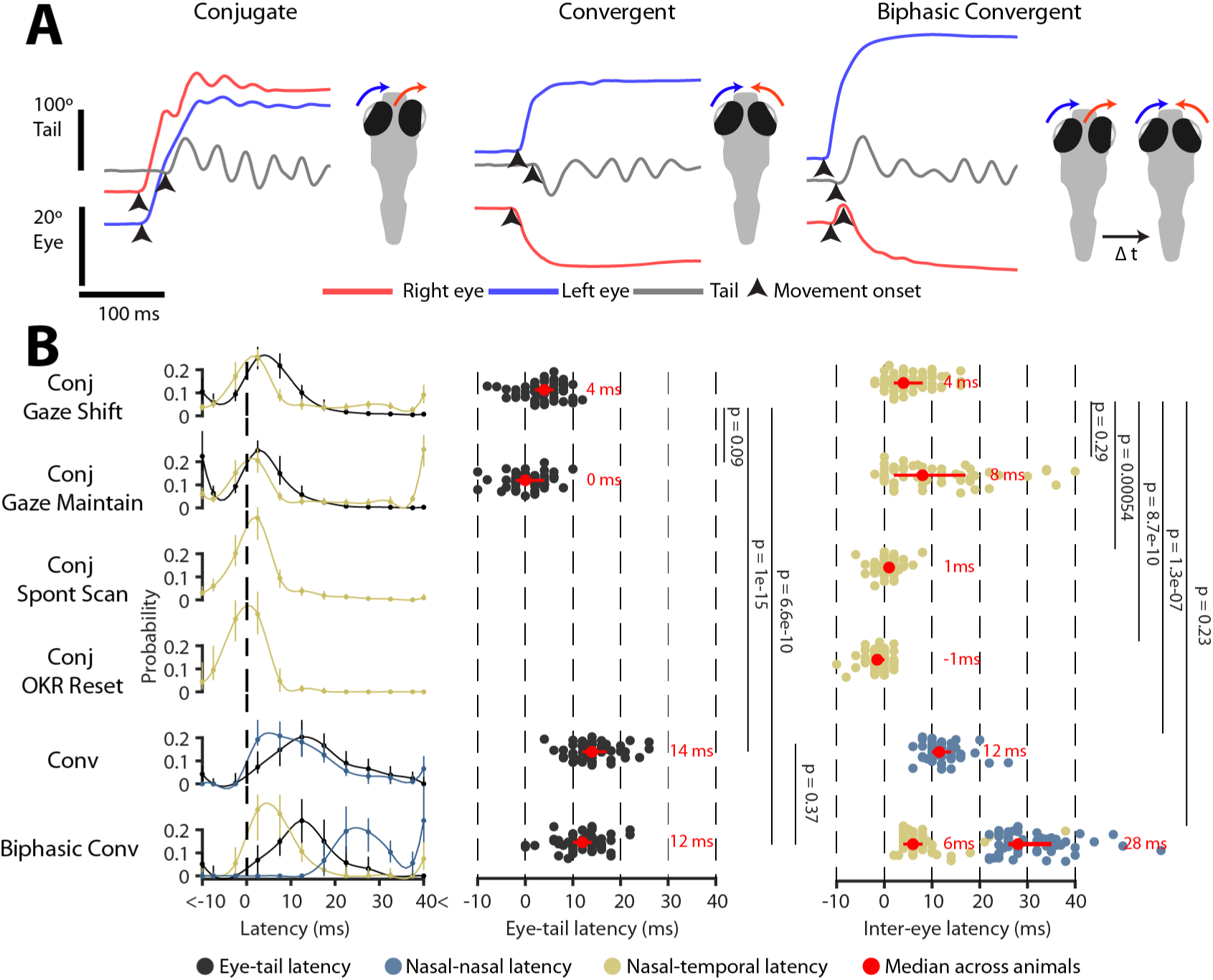
Timing relationships between eye and tail movements vary across saccade types. (A) Estimation of movement onset times (arrowheads) for the eyes and tail for exemplar conjugate, convergent and biphasic convergent saccades. (B) *Left:* Distribution of eye-tail and inter-ocular latencies across saccade types. All latencies measured relative to first nasal eye rotation. Median ± IQR proportions across *N* = 58 animals with spline fits. *Middle, right:* Median eye-tail (middle) and inter-ocular (right) latencies per animal. Median *±* IQR across animals in red with p-values from Kruskal-Wallis with Dunn-Sidak post-hoc tests. (Note that average latency values in main text are mean *±* SEM).

The time interval between movement initiation of the two eyes and the eyes and tail differed across saccade types [Figure 5B]. For conjugate saccades occurring in the absence of tail movements (scanning saccades in stationary animals and OKR fast phases), the latency between eye movements was small (*≤* 5 ms in 58.1 *±* 2.0% cases, *N* = 58 animals), with nasal (adducting) eye movement tending to occur shortly before temporal (abducting) eye movement (mean inter-ocular latency 1.1 *±* 0.5 ms, *p* = 0.016 t-test versus zero). However, when conjugate saccades were accompanied by swims, inter-ocular latencies were substantially longer and tail movement was coincident with the first (nasal) eye rotation (inter-ocular latency 14.2 *±* 0.7 ms, *p* = 2.3 *×* 10^-26^; nasal-tail latency 1.1 *±* 0.6 ms, *p* = 0.052; t-tests vs zero). We did not observe a significant difference in these timing relationships when comparing small conjugate saccades that maintain gaze direction during forward locomotion versus saccades that shift gaze in coordination with body turns [Figure 5B].

Biphasic convergent saccades are defined by an initial conjugate rotation followed by the abducting eye reversing direction and moving nasally (right eye in example in [Figure 5A]). By analysing movement timing, we found that the initial conjugate movement had inter-ocular latency of 13.4 *±* 1.0 ms, which was not significantly different to conjugate gaze-shifting saccades [Figure 5B]. The second nasal rotation then followed with a long delay (36.2 *±* 1.1 ms). By contrast, regular convergent saccades showed significantly longer inter-ocular and eye-tail latency as compared to conjugate saccades (mean inter-ocular latency 15.6 *±* 0.5 ms; eye-tail latency 13.8 *±* 0.6 ms).

In sum, saccadic eye movements of larval zebrafish are coordinated with body movements on a millisecond timescale. As observed in other species, latencies are incompatible with sensory feedback and instead indicate that eye and body movements are controlled by a common neural command. However, timing relationships differ across saccade types, implying that distinct patterns of circuit activity coordinate the two eyes and in particular the eyes and body for convergent versus conjugate saccades.

### Velocity profiles and main sequence relationships indicate that distinct extraocular motoneuron activity controls different saccade types

To produce a saccade, extraocular motoneurons generate a stereotypical pulse-glide-step firing profile in which a burst of high-frequency spiking (pulse) first accelerates the eye and firing rate then decays (glide) to a lower and sustained rate (step) that holds the eye in its new position against centripetal elastic forces (Sparks, 2002). Due to the regular properties of the ocular plant, features of motoneuron activity can be readily inferred from the kinematics of the eye movement (Bahill & Troost, 1979; Robinson, 2022a,c). We therefore used high temporal resolution tracking data to assess the kinematics of conjugate and convergent saccades to estimate if they might be produced by distinct brainstem motor control signals (*N* = 58 tethered larvae).

Conjugate and convergent saccades had distinct eye position and velocity time courses. When comparing the adducting (nasal) eye movements that are common to both, we observed that for saccades *>* 15*^◦^*, the eye reached its final position more quickly for convergent saccades, whereas conjugate saccades took longer to obtain final eye position [Figure 6A]. Convergent saccades had rather symmetrical velocity profiles and peak velocity progressively increased with saccade amplitude [Figure 6B]. In contrast, peak velocity was substantially lower for large conjugate saccades and velocity profiles were markedly asymmetric, declining slowly with a protracted tail. These features indicate that conjugate saccades are hypometric, showing a dynamic undershoot and then slowly obtaining final position. We considered the possibility these differences might be explained by systematic differences in starting eye position in the orbit; however, when we binned saccades by starting position, the same hypometric pattern for conjugate saccades was observed [Figure S4A]. From these observations we can infer that for large conjugate saccades, the ‘pulse’ on medial rectus motoneurons is poorly matched to the required change in eye position.

**Figure 6:**
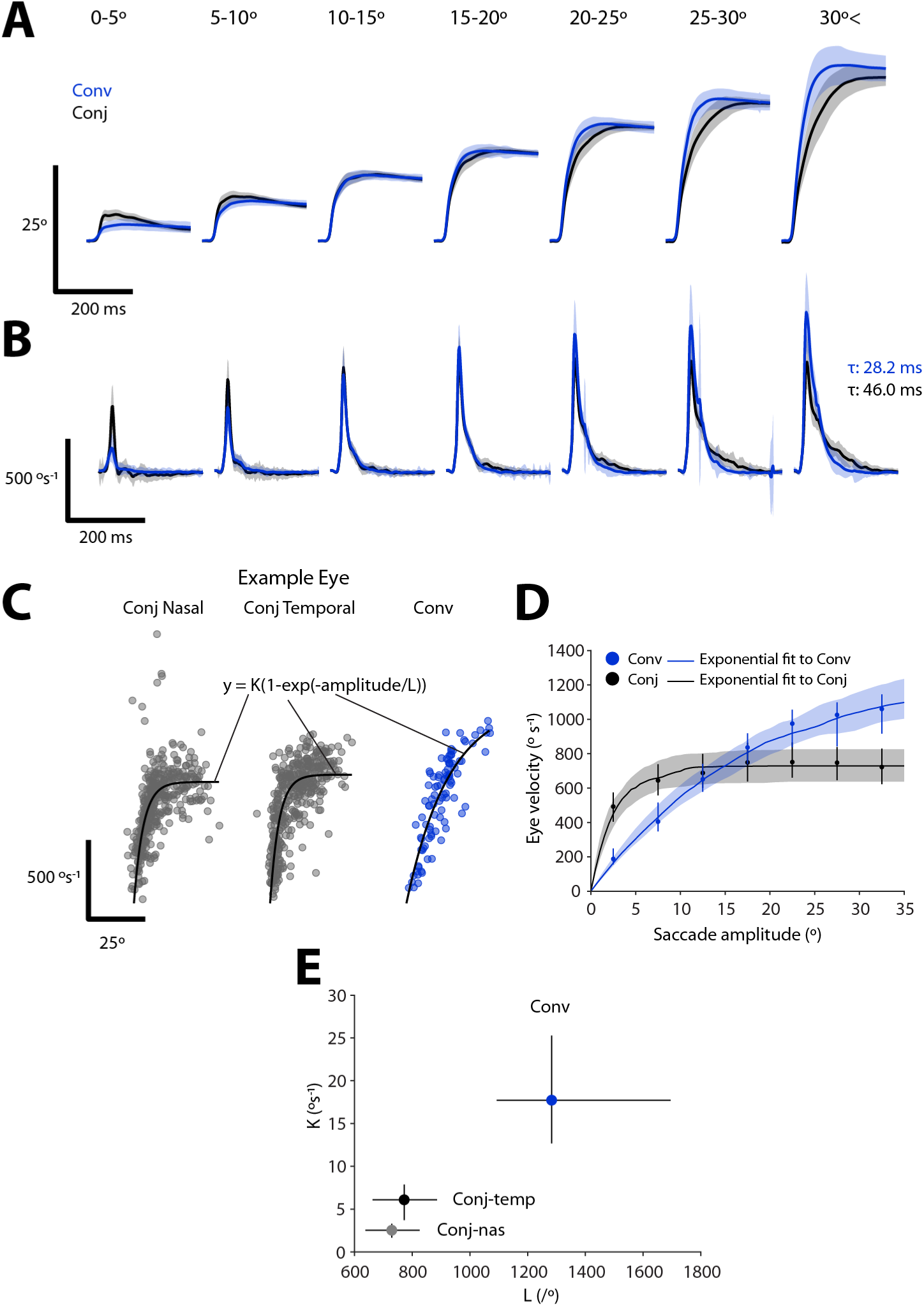
Distinct velocity main sequence relationships for convergent and conjugate saccades. (A-B) Eye position (A) and velocity (B) time series for adducting (nasal) saccadic eye movements from convergent and conjugate saccades of different amplitudes. Mean ± SD for *N* = 116 eyes from 58 fish. (C) Velocity main sequence relationship for one example eye with exponential fits. (D) Average velocity main sequence for convergent and conjugate nasal saccades. Lines and shading show median *±* IQR across exponential fits to *N* = 116 eyes from 58 fish. Points indicate median (*±* IQR) eye velocity binned by amplitude. (E) Fit coefficients for saccade types (median *±* IQR). *See also Figure S4*.

A defining feature of saccadic eye movements is the ‘main sequence’ relationship, wherein eye velocity increases as a function of saccade amplitude before saturating (Bahill et al., 1975c). To examine this relationship, we fit exponential functions (Baloh et al., 1975; Gibaldi & Sabatini, 2021) to the velocity and amplitude of saccades of individual eyes [Figure 6C], and then computed average main sequence relationships [Figure 6D]. This revealed that the adducting eye follows distinct main sequence relationships during conjugate versus convergent saccades ([Figure 6E], *AIC* = 97.8 *±* 0.15*, p* = 1.0 *×* 10*^−^*^28^*, N* = 83 eyes). Specifically, while small conjugate saccades are faster than similarly sized convergent saccades, conjugate saccade velocity saturates rapidly, reaching a plateau of *∼* 700*^◦^*/s above *∼* 15*^◦^*. Notably, both gaze-maintaining and gaze-shifting conjugate saccades followed the same main sequence pattern [Figure S4B], supporting the idea that the former are true saccadic eye movements and that all conjugate saccades are generated by a common underlying pattern of neural circuit activity. By contrast, for convergent saccades, velocity increased as a more linear function of saccade amplitude [Figure 6D]. These results reveal different patterns of medial rectus motoneuron population activity for the two saccade types. Specifically, the pulse component saturates for conjugate saccades, such that peak velocity fails to keep pace with amplitude, but continues to increase for large amplitude convergent saccades.

We also analysed biphasic convergent saccades to assess if all three component eye movements were true saccades and if so, whether they were generated by neural control signals similar to convergent or conjugate saccades [Figure S4C]. We observed that the first nasal eye movement followed the same main sequence relationship as we had observed for regular convergent saccades [Figure S4D left]. By contrast, the temporal movement of the opposite eye (mean amplitude 4.5*±*0.1*^◦^*) followed a velocity main sequence comparable to conjugate saccades [Figure S4D right]. This was also supported by linear (rather than exponential) fits of saccade velocity versus amplitude, which showed comparable slopes for convergent saccades and the first nasal component of biphasic saccades (slope *∼* 45*/s*), whereas the first temporal movement had a much greater slope, equivalent to small amplitude conjugate saccades (*∼* 90*/s*) [Figure S4E]. The second nasal movement followed a main sequence with lower velocity as compared to regular convergent saccades, likely a result of the immediately preceding temporal rotation. Altogether, this analysis indicates that biphasic convergent saccades comprise three saccadic eye movement components that are likely to be controlled by monocular premotor commands (see Discussion).

## Discussion

By analysing thousands of rapid eye movements in tethered and freely swimming zebrafish larvae, we identified five saccade types that differ in oculomotor kinematics and binocular coordination and which are used in distinct behavioural contexts. We defined four roles for conjugate saccades and found that they are coordinated with swims of either the same, or opposite, laterality to shift or maintain gaze, respectively, during locomotion. By contrast, convergent saccades play a specialised role in generating precise, goal-directed gaze shifts that enable zebrafish to foveate their prey from the onset of hunting sequences. Conjugate and convergent saccades differed in the precise timing of binocular and eye-body coordination and followed different velocity main sequence relationships, pointing to differences in underlying physiological control. Our work aligns with recent efforts to characterise active vision during naturalistic behaviour (Meyer et al., 2020; Michaiel et al., 2020) and complements recent studies in zebrafish that have comprehensively defined the animal’s locomotor repertoire (Marques et al., 2018) and determined how swims are selected and sequenced during exploration and hunting (Dunn et al., 2016b; Wolf et al., 2017; Bolton et al., 2019; Mearns et al., 2020; Johnson et al., 2020). By uncovering how and when saccades are used to redirect gaze, this study provides insight into the visuomotor strategies that organise behaviour and will guide experiments examining circuit control of eye movements, eye-body coordination and visuomotor processing. Finally, we note that our estimate of the zebrafish saccadic repertoire may be incomplete. We assayed only a subset of (visually guided) behaviours and restricted our analysis to horizontal eye movements. Analysis of a broader range of behaviours and tracking of vertical and torsional eye movements (Bianco et al., 2012) may reveal additional types of, or uses for, saccades, perhaps in coordination with pitch/roll postural adjustments (Ehrlich & Schoppik, 2019).

### Conjugate saccades and visual exploration

Zebrafish used conjugate saccades to recentre the eyes (i.e. fast phases) during the optokinetic nystagmus, to redirect gaze without an accompanying head/body rotation, and in coordination with swims to either shift or maintain gaze direction. In our classification, conjugate saccades were characterised by both eyes rotating in the same (clockwise or counterclockwise) sense, but the amplitude of left and right eye movements were not necessarily equal. Dissimilar amplitudes were clearly observed for the large conjugate saccades of stationary animals, where the abducting saccade was typically greater. By producing a slight divergence, these saccades expand the visual field, compatible with the idea that they allow the animal to survey a broad region of its environment even when at rest. These scanning saccades will reduce the (time-averaged) extent of the blind spot in the visual field behind the animal (Bianco et al., 2011) and may be part of an active sensing strategy, for example to sample luminance gradients (Wolf et al., 2017). An alternative, non-mutually exclusively hypothesis, is that these saccades help overcome visual adaptation (Samonds et al., 2018). In any case, it is clear that like other fish species (Harris, 1965; Hermann & Constantine, 1971; Easter, 1971), zebrafish use saccadic eye movements alone to redirect gaze.

Conjugate saccades are coordinated with head/body movements in both foveate and afoveate species (the ‘afoveate saccadic system’, Robinson (2022b)) and accordingly, we found that larval zebrafish head/body reorientations exceeding *∼* 10*^◦^* were paired with conjugate saccades of the same laterality. In contrast to primates (Freedman, 2008), gaze shifts of increasing amplitude were not accomplished by systematically varying the relative contributions of the eyes versus head/body. Instead, body reorientation scaled linearly with the overall gaze shift and conjugate eye amplitude was quite variable. However, as in other species, post-saccadic eye position was displaced in the direction of locomotion. By contrast, forward swims were coincident with conjugate saccades of the opposite laterality. This was surprising, because, to our knowledge, saccades paired with head/body movements of opposite directionality have not been described previously. We believe that these are bona fide saccadic eye movements as they conform to the velocity main sequence and precede the spino-ocular coupling reflex that compensates for the head yaw during swimming (Straka et al., 2022). By opposing body reorientation, these saccades help to maintain the animal’s line of sight during forward locomotion and so we refer to them as gaze-maintaining. Again, the animal ‘looks where it is going’. Notably, these saccades have an amplitude of approximately half of the body reorientation. While at first glance this might appear insufficient, as noted by Easter & Johns (1974), ‘perfect’ compensation would stabilise a visual plane at infinity, which is likely of little use in aquatic environments, whereas partial compensation stabilises the visual world more proximal to the animal. Considering that a typical forward swim bout produces *∼* 1.2 mm displacement (Severi et al., 2014), we estimate that gaze-maintain saccades stabilise a visual plane *∼* 24 mm from the head. Calcium imaging experiments have identified activity in rhombomeres 2 and 3 correlated with the direction of tail and eye movement (Dunn et al., 2016b; Wolf et al., 2017; Ramirez & Aksay, 2021), but because eye and body rotations were highly correlated in these assays, it is unclear if this brain region controls either or both motor outputs. The switch in saccade directional contingency described here creates a clear distinction between forward swims and turns and provides a handle for future studies to dissect the neural commands that generate coordinated eye-body motor programmes.

### Convergent saccades, binocular vision and foveation of prey

Convergent saccades are a defining feature of larval zebrafish hunting behaviour (Bianco et al., 2011; Trivedi & Bollmann, 2013; Bianco & Engert, 2015). Both regular and biphasic convergent saccades obtained eye velocities in excess of 600*^◦^*/s; these eye movements are therefore unlike the slow (*∼* 30*^◦^*/s) fusional vergence movements of primates and instead more similar to primate disjunctive saccades, which are used to rapidly shift fixation between points at different distances and directions in three-dimensional space (Enright, 1984; Quinet et al., 2020). Zebrafish use these specialised saccadic eye movements to engage a predatory mode of gaze during hunting and a high vergence angle is then sustained throughout prey-tracking until after the final capture strike. Immediately prior to capture, the eyes are symmetrically and maximally converged such that the most proximal point of binocular overlap is directly ahead of the larva and only 400 *µ*m from the midpoint of the eyes (Bianco et al., 2011). On the basis of these observations, we proposed that eye convergence likely supports a simple stereopsis mechanism allowing larvae to estimate that prey is located at a specific point in egocentric space (the ‘strike zone’) and release a capture swim, a hypothesis that has received support from elegant lens-removal experiments (Mearns et al., 2020). However, because convergent saccades occur at the onset of hunting and are lateralised towards prey (Trivedi & Bollmann, 2013; Henriques et al., 2019), it seems likely that binocular vision plays additional roles throughout prey-tracking. Here we find that zebrafish smoothly control the conjugate (version) component of convergent saccades in accordance with retinotopic prey azimuth. By analysing prey position in a gaze-referenced coordinate space that is relevant for visual perception, we found that the combined eye-body gaze shift redirects the binocular visual field towards prey with surprisingly high gain (*∼* 0.9). The effect of this ‘visual grasp’ is to bring prey images to the high acuity area (‘fovea’) of the retina, which contains an elevated density of UV cones and additional physiological specialisations for detection of UV-bright prey (Yoshimatsu et al., 2020). This suggests that zebrafish may use convergent saccades as part of a visuomotor strategy to achieve high signal-to-noise detection of low-contrast prey objects. Moreover, in half of trials, the first orienting manoeuvre during hunting was sufficient to bring prey images to the HAA of both eyes. Beyond further increasing detection sensitivity (by a factor of up to √2), it seems plausible that binocular foveation, along with an internal (efference copy) estimate of eye position, allows larvae to estimate prey distance by a simple algorithm equivalent to triangulation. In support of this idea, Bolton et al. (2019) has shown that swim vigor is modulated by prey distance during prey-tracking (at distances *<* 4 mm), indicating that larvae have the means to estimate this variable. Whereas conjugate saccades were near-coincident with tail movements, convergent saccades led the tail by 14 ms. It has been suggested that moving the eyes first helps to compensate for the sluggishness of visual processing during gaze shifts (Robinson, 2022b); such a function would therefore imply a particular importance for maintaining prey perception during hunting.

### Saccade type-specific neural control

A key finding of our study is that different saccade types display distinct kinematics, velocity main sequence relationships, and binocular and eye-tail timing relationships, together suggesting differences in underlying neural control. Modulation of saccade latency and kinematics has been shown in several contexts. For example, in primates, saccades in the dark are consistently slower than those in the light, reactive saccades are slightly faster than voluntary saccades, decision making tasks can influence saccade velocity, and the shape of velocity profiles is modulated depending on whether or not a head movement contributes to gaze shifting (Freedman, 2008; Gremmler & Lappe, 2017; Seideman et al., 2018; Robinson, 2022d). Our findings show that in zebrafish larvae, eye movement kinematics vary systematically across saccade types, where those types are defined by different patterns of binocular coordination and have distinct ethological roles. A velocity main sequence has previously been described for spontaneous conjugate saccades and OKR fast phases in larval zebrafish (Chen et al., 2016; Leyden et al., 2021) and our results extend these findings to show that all subtypes of conjugate saccade (including gaze-shift and gaze-maintain saccades paired with swims) conform to the same main sequence relationship. This in turn suggests that all conjugate saccades are controlled by the same peripheral circuits, in line with established ideas about saccade generation (Bahill et al., 1975c; Robinson, 2022d). This velocity main sequence showed an inflection point at *∼* 15*^◦^*, similar to saccades in other species including goldfish (Salas et al., 1997) and humans (Gibaldi & Sabatini, 2021). At greater amplitudes, there is minimal further increase in peak eye velocity, which indicates that the phasic ‘pulse’ component of extraocular motoneuron activity reaches a ceiling and is no longer able to scale with the amplitude of the gaze shift. This is concordant with the pronounced asymmetry we observed in the velocity time course, where after an initial rapid acceleration, hypometric conjugate saccades obtained final eye position with a much slower, ‘glissadic’ eye movement, likely during the slide and/or step phase of motoneuron firing. Here it is pertinent to note recent work that has revealed a very broad range of time constants in extraocular motoneuron activity matched to a similarly broad range of viscoelastic time constants in the oculomotor plant (Miri et al., 2022). In future, it would be interesting to apply these models to the various types of saccadic eye movement we describe to better estimate underlying neural activity. Convergent saccades had quite different position/velocity kinematics and main sequence relationships. While small amplitude (*<* 10*^◦^*) saccades were slower than equivalent conjugate saccades, the main sequence showed substantially less saturation with peak velocity continuing to increase across the full dynamic range of saccade amplitude (*∼* 35*^◦^*). Accordingly, large convergent saccades did not appear hypometric and had more symmetric velocity time courses indicating that the pulse on medial rectus motoneurons was matched to the required change in eye position.

What might be the physiological basis for these kinematic differences? One possibility is that there is a distinct (or additional) population of medial rectus motoneurons responsible for nasal eye rotations during convergent saccades. However, although it has been long debated, there is currently little if any evidence for groups of motoneurons with specialised roles in particular types of eye movement, let alone specific subtypes of saccade. Although there are two major types of extraocular motoneuron, with distinct molecular properties, patterns of afferent input and synapse termination on extraocular muscle fibres, physiological data reveals a smooth continuum of functional properties and it is generally assumed that all extraocular motoneurons participate in all classes of eye movement (Evinger & Baker, 1991; Hernández et al., 2019; Horn & Straka, 2021). Nonetheless, extraocular motoneurons do show substantial variation in recruitment threshold and position and velocity sensitivity. Therefore differences in afferent input may give rise to distinct patterns of motoneuron recruitment and activity to bring about saccadic eye movements with type-specific kinematics. In future work, neural activity recordings in the zebrafish brainstem will be a powerful tool to discover saccade type-specific neural populations and evaluate motoneuron population activity as a function of both saccade type and oculomotor kinematics.

### Biphasic convergent saccades

Finally, the unusual properties of biphasic convergent saccades warrant special mention. These comprise three closely coordinated saccadic movements; an initial conjugate eye rotation, shortly followed by reversal of the abducting eye. Closely spaced saccades have been described in humans (Bahill et al., 1975a) and the reversal in biphasic convergent saccades is perhaps most reminiscent of dynamic overshoot, a common phenomenon in which a primary saccade is immediately followed by a small secondary saccade in the opposite direction (Bahill et al., 1975b; Kapoula et al., 1986). Dynamic overshoot is also a monocular phenomenon, typically of the abducting eye, and the return eye movement has saccadic velocity. It has been suggested that a braking pulse of neural activity normally functions to bring the eye to rest at the end of a saccade and dynamic overshoot may occur when this pulse is excessively large (Kapoula et al., 1986). Along similar lines, biphasic convergent saccades might arise due to errors in the amplitude and/or timing of multiple saccadic commands that normally control (routine) convergent saccades. The nature of the premotor commands that control binocular eye movements has long been debated (King, 2011; Coubard, 2013): Hering’s Law posits that both eyes receive identical (conjugate and vergence) neural commands, whereas Helmholtz argued for independent control of each eye. The fact that the first nasal and temporal eye movements of biphasic convergent saccades conform to different main sequence relationships (characteristic of convergent and conjugate saccades respectively), seems compatible with emerging evidence in favour of a monocular control framework (Zhou & King, 1998; King & Zhou, 2000; Sylvestre et al., 2003; Cullen & Van Horn, 2011).

## Acknowledgements

The authors thank members of our lab for helpful discussions and critical feedback on the project and the UCL Fish Facility staff for fish care and husbandry. Funding was provided by a Sir Henry Dale Fellowship from the Royal Society & Wellcome Trust (101195/Z/13/Z) and a Senior Research Fellowship from the Wellcome Trust (220273/Z/20/Z) to I.H.B., and a Wellcome Trust 4 year Neuroscience PhD studentship to C.K.D.

## Author Contributions

Conceptualisation and Methodology: C.K.D. and I.H.B.; Investigation: C.K.D., J.Y.N.L.; Analysis: C.K.D.; Writing: C.K.D. and I.H.B.; Supervision and Funding Acquisition: I.H.B.

## Methods

### Zebrafish

Zebrafish lines were maintained in the Tübingen background. Larvae were reared in fish-facility water on a 14/10 h light/dark cycle at 28.5*^◦^*C and were fed *Paramecia* from 4 dpf onwards. All larvae carried the *mitfa* (Lister et al., 1999) skin-pigmentation mutation. The 152 animals used for tethered behavioural analysis carried transgenes as follows: 76 animals carried Tg(*elavl3*:H2B-GCaMP6s)jf5Tg (Freeman et al., 2014). 60 animals carried Tg(*pvalb6*:KalTA4)u508 (Antinucci et al., 2019) and Tg(UAS:GCaMP6f)icm06 (Böhm et al., 2016). 4 animals carried Tg(*isl1*:GFP)rw0Tg (Higashijima et al., 2000) and Tg(*elavl3*:H2B-GCaMP6s)jf5Tg. 6 animals carried Tg(*vsx2*:Gal4FF)nns18Tg (Kimura et al., 2013), Tg(UAS:RFP) and Tg(*elavl3*:H2B-GCaMP6s)jf5Tg. 6 animals carried Tg(*elavl3*:GCaMP7f)u343Tg. Eight animals were used for free-swimming assays analysis and were not transgenic. The sex of the larvae is not defined at the early stages of development used for these studies. Experimental procedures were approved by the UK Home Office under the Animals (Scientific Procedures) Act 1986.

### Behavioural tracking in tethered larvae

Larvae were tethered in 3% low-melting point agarose gel in a 35 mm petri dish lid and sections of gel were carefully removed using an opthalmic scalpel to allow free movement of the eyes and tail below the swim bladder. Larvae were allowed to recover overnight before testing at 6 or 7 dpf. Behavior was tracked whilst animals underwent two-photon calcium imaging using a custom microscope as described in (Antinucci et al., 2019). Eye movements were tracked at either 60 or 300 Hz using a FL3-U3-13Y3M-C camera (Point Grey) that imaged through the microscope objective. Tail movements were imaged at 420 Hz under 850 nm illumination using a sub-stage GS3-U3-41C6NIR-C camera (Point Grey). Horizontal eye position and tail posture (defined by 13 equidistant x-y coordinates along the anterior-posterior axis) were extracted online using machine vision algorithms (Bianco & Engert, 2015).

Two projectors were used to present visual stimuli. The first (Optoma ML750ST) back-projected stimuli onto a curved screen placed in front of the animal at a viewing distance of 35 mm while the second (AAXA P2 Jr) projected images onto a diffusive screen directly beneath the chamber. Wratten filters (Kodak, no. 29) were placed in front of both projectors. Visual stimuli were designed in MATLAB using Psychophysics Toolbox (Brainard, 1997). Prey-like moving spots comprised 6*^◦^* or 12*^◦^* bright or dark spots (Weber contrast +1 or -1 respectively) moving at 30*^◦^*/s either left*→*right or right*→*left across 152*^◦^* of frontal visual space. For dark flashes, both projectors were switched to zero pixel value for 3 s. Looming stimuli comprised expanding dark spots (Weber contrast -1) that simulated an object approaching at constant velocity (10*^◦^*–70*^◦^*, L/V 490 ms) (Sun & Frost, 1998). Optomotor stimuli comprised drifting sinusoidal gratings (wavelength 10 mm, velocity 10 mm/s, Michelson contrast 1) presented from below and moving in four cardinal directions with respect to the animal. Optokinetic stimuli comprised drifting sinusoidal gratings (wavelength 19*^◦^*, velocity 0.3 cycles/s, Michelson contrast 0.5) presented in front of the animal and moved left-to-right and right-to-left. For all experiments, stimuli were presented in a pseudo-random sequence with 30 s inter-stimulus interval.

Microscope control, stimulus presentation and behaviour tracking were implemented using LabView (National Instruments) and MATLAB (MathWorks).

### Behavioural tracking in freely swimming larvae

Free-swimming behaviour was recorded in a similar manner to Henriques et al. (2019). In brief, behaviour was recorded in a 35 mm petri dish with 3% low-melting point agarose placed along the walls to limit thigmotaxis. The chamber was placed on a horizontal platform onto which visual stimuli could be presented (Acer C202i projector) via a cold mirror from below. Images were acquired at 300 Hz under 850 nm illumination using a Mikrotron EoSens 4CXP camera equipped with a machine vision lens (Kowa) and a 850 nm bandpass filter.

Visual stimuli were designed in MATLAB using Psychophysics toolbox. Optomotor stimuli comprised sinusoidal gratings (wavelength 8 mm, velocity 8 mm/s, Michelson contrast 1, duration 6 s) that drifted at 90*^◦^* to the left or right with respect to the fish, with stimulus direction locked to fish orientation and updated in real-time. Stimuli were presented with a minimum interstimulus interval of 120 s and only when the centroid of the larva was within a predefined central region (*∼* 11 mm from arena edge). If the fish strayed out of this region, a concentric grating was presented that drifted towards the centre of the arena to attract the fish back. Only behaviour data from within this central region was analysed to avoid tracking errors caused by reflections from the chamber edge.

At the beginning of the experiment 10-30 *Paramecia* were added to the dish to promote hunting behaviour.

Eye and tail kinematics were tracked online as described in Henriques et al. (2019). Throughout the experiment a cropped (23.9 mm *×* 23.9 mm, 13.0 mm/px) movie centred on the centroid of the larva was recorded to allow subsequent analysis of hunting orientations (see below). Each experiment lasted around 45 min. Camera control, online tracking and stimulus presentation were implemented using LabVIEW (National Instruments) and MATLAB (Mathworks).

### Saccade detection and classification

Raw eye position traces were interpolated onto a 100 Hz time-base and low-pass filtered with a cut-off frequency of 1 Hz. Rapid eye movement events were detected as peaks in the convolution of filtered eye position with a step function (width 160 ms), with the timepoint of the peak providing a first coarse estimate of movement initiation time. Rapid eye movement events of the left and right eye that occurred within 100 ms of one another were paired and treated as a single binocular event. After this pairing step, events that occurred within 300 ms of a preceding event were discarded, to limit overlap between windows for calculating saccade metrics (see below) and because manual inspection revealed that these movements were rarely saccadic.

To reliably estimate eye position and velocity metrics, raw eye position traces were interpolated onto a 500 Hz timebase and smoothed with a custom LOWESS function. A more refined estimate of onset time was determined by convolving smoothed eye position with two step functions of width 100 ms and 40 ms, taking the product between both convolutions and thresholding the output within a 400 ms window spanning the initial estimate of saccade time. The custom LOWESS function was designed to reduce noise in eye position traces without flattening changes in eye position during saccades. This involved applying the MATLAB lowess function with two different spans depending on whether a saccade-like change in eye position was detected. A shorter span was used during stepwise changes in eye position. For free swim data this span was 80 ms. For tethered data no smoothing was done. A larger span was used outside of stepwise changes. This was 133 ms for free swim and 33 ms for tethered data. Stepwise changes were defined by convolving raw eye position traces with a step function and thresholding, in a similar manner to the rapid eye movement detection procedure.

For each rapid eye movement event we evaluated: (a) pre-saccadic eye position, as median eye position during a 200 ms window immediately prior to onset time; (b) max post-saccadic eye position, as the eye position within a 200 ms window starting at onset time that had the greatest absolute deviation from eye position at onset time; (c) median post-saccadic eye position, as median eye position over a 200 ms window starting at the timepoint corresponding to max post-saccadic eye position; (d) eye velocity (cw and ccw), as the maxima and minima, respectively, of the time derivative of eye position, determined by the MATLAB gradient function over a 150 ms window centred at onset time. We then used these measures to calculate nine oculomotor metrics describing each (binocular) rapid eye movement event: *Amplitude* (left and right eye), was the difference between median post-saccadic eye position and pre-saccadic eye position. *Max-median amplitude* (left and right eye), was the difference between max post-saccadic eye position and median post-saccadic eye position and quantifies the degree to which eye position is maintained following a saccade. *Velocity* (cw and ccw for both left and right eye), as described above. *Vergence* was the difference between median post-saccadic eye position of the right and left eye.

To examine variation in oculomotor kinematics and categorise rapid eye movements, we embedded eye movement events in a low dimensional space and applied a density based clustering procedure. To do this, data from each animal was first winsorized (0.5-99.5th percentile) and z-scored. Our initial embedding and clustering was performed using data from tethered larvae, excluding events that initiated during a swimming bout (213,462 of 335,442 events (63.6%) from *N* = 152 fish). This helped to ensure we used high quality tracking data without artefacts caused by swim-induced changes in eye/head position. Datapoints were embedded into two dimensions using a MATLAB implementation (Meehan et al., 2022) of UMAP (McInnes et al., 2018) (run_umap, metric=Euclidean, min_dist=0.11, n_neighbours=199) and the output clustered using DBSCAN (Ester et al., 1996), with epsilon = 0.34 and minimum point threshold = 570. Un-clustered points within 3 units of UMAP space to a cluster edge were assigned to a cluster within this radius; the event was assigned to the cluster that had the most successive increases in point density binned along a straight line connecting the event and the cluster centroid. Supervised embedding, using the previous UMAP solution as a template, was applied to tethered events that initiated during a swim bout (121,980 events) as well as data from free-swimming larvae (10,569 from 8 fish). These datapoints were assigned the most common cluster identity from 100 nearest neighbours in the embedding space; however, if those 100 nearest neighbors were separated from the target event by a median Euclidian distance exceeding 0.3 in UMAP space (1,396 tethered events, 0 free-swimming events) then no identity was assigned.

Following this initial clustering we observed that some biphasic convergent saccades were assigned to the Conv cluster, rather than the two BConv clusters. We therefore implemented an additional procedure to detect and reassign BConv events. Specifically, biphasic convergent saccades were defined by having one eye that moved in a temporal direction with velocity and eye displacement exceeding thresholds: The velocity threshold was 60*^◦^*/s for tethered data and 40*^◦^*/s for free-swimming data. The eye displacement threshold was one standard deviation of eye position over a 150 ms window terminating 100 ms prior to onset time.

### Swim kinematic analysis

Raw tail tracking data from tethered larvae comprised 13 x-y centroids defining the midline of the tail. Consecutive centroids define 12 tail segments and vectors of 11 inter-segment angles were computed for each timepoint. Raw tail tracking data from freely swimming larvae comprised 9 x-y centroids, producing 7 inter-segment angles that were interpolated to 11 inter-segment angles to maintain consistency across datasets. Matrices of inter-segment angles over time were interpolated onto a uniform 1000 Hz timebase and smoothed in 2D using a 2-segment-by-7-ms filter. Next, we computed the cumulative sum of inter-segment angles, *γ*, which was filtered (MATLAB sgolayfilt, order=3, framelength=9) and median subtracted. Thus, changes in tail posture are represented as the time-varying cumulative bend angle along the anterior-posterior axis of the tail: *γ_s,t_*, for cumulative inter-segment angle *s* at time-point *t*. To identify swim bouts, we first estimated tail angular velocity, *v_t_* by differentiating *γ*_11_*_,t_*, taking its absolute value and filtering (40 ms box-car). We also computed the envelope, *f_t_*, as the maximum absolute value of *γ*_11_*_,t_* within a 9 ms sliding window. The start of swim bouts were identified at time-points where *v_t_ >* 800 deg/s and *f_t_ >* 7 deg, and the end of swim bouts was defined when *v_t_ <* 200 deg/s and *f_t_ <* 10 deg. Bouts less than 61 ms in duration were excluded.

We identified individual halfbeats (leftwards and rightwards excursions of the tail) by finding the maxima and minima of *γ*_9_*_,t_*. For tethered larvae we used the sign of the first half-beat for cumulative inter-segment angle 11, *θ*_11_,_1_*_st_*, to define swim direction (left/right) and its amplitude as a proxy for swim lateralisation. For free-swimming larvae, we computed the change in body orientation, Δ*_ori_* by taking the difference between body orientation 50 ms before and 7.5 ms after a swimming bout; the sign of Δ*_ori_* defined swim direction.

Rapid eye movements were considered coincident with a swim if their onset time was from 200 ms prior to swim initiation to 200 ms after swim termination.

### Contextual deployment of saccades

In Figure 2, contexts were defined based upon swim types and visual stimuli and the frequency of saccade types occurring in these contexts was evaluated. The contexts, which are not mutually exclusive, were defined as follows: Swim L/R, swims for which *|*Δ*_ori_| ≥* 10*^◦^* to left/right for free-swimming or *|θ*_11,1_*_st_| ≥* 25*^◦^* for tethered fish. Swim F, were swims with *|*Δ*_ori_|* (or *|θ*_11,1_*_st_|*) below these thresholds. OKR-L/R, optokinetic gratings drifted to the left/right and there was no accompanying swim bout. Prey spot, presentation of prey-like moving spot stimuli. Orient to spot, first saccade during presentation of prey-like moving spot accompanied by swim with direction corresponding to spot laterality. Loom <30 deg and Loom > 30 deg, looming stimulus subtended visual angle less than and greater than 30°, respectively. Dark flash, dark flash stimuli. No-stim-no-swim, inter-stimulus intervals and no coincident swim bout.

### Orientating responses to prey in freely swimming larvae

The initiation of a hunting sequence was defined as a convergent saccade that increased vergence above a threshold. This was determined for each animal by fitting two Gaussians to the bimodal distribution of vergence angles measured across the experiment and setting the threshold to one standard deviation below the centre of the higher Gaussian. Next, for each convergent saccade, five consecutive imaging frames (starting 10 ms prior to saccade onset) were assessed for putative prey targets. Putative targets were identified by Gaussian filtering, thresholding and detecting small binary objects (148 *< Area <* 889 *µm*^2^ *∩* 385 *< Length <* 1540 *µm*). If at least one putative target was identified across the five images, target positions were determined manually using a custom MATLAB GUI. Instances where there were multiple prey objects in the animal’s visual field (see below) were not assessed to avoid ambiguity in target identification.

Orientations to prey targets were decomposed into three components: pYaw, was the change in prey angular position attributable to the change in orientation of the larva during the swim bout coincident with the convergent saccade. This was equal to Δ*_ori_* for said swim bout. pTrans, was the change in prey angular position attributable to translation of the animal’s head during the swim bout coincident with the convergent saccade. This was computed as *α_post_ −α_pre_*, where *α_pre_* was the angle between the vector connecting the midpoint of the eyes and the prey target and the vector defining head orientation at the time of convergent saccade initiation. *α_post_* was calculated 7.5 ms after completion of the swim bout as the angle between the vector connecting the midpoint of the eyes and the prey target and the vector defining head orientation *at the previous time of convergent saccade initiation*. In this way, the effect of body orientation change was eliminated. pConv, was the change in the angle of the vector connecting the midpoint of the eyes to the most proximal point of binocular overlap before versus after the convergent saccade. The most proximal point of binocular overlap was defined as the point at which the nasal limits of the left and right visual fields overlapped, with each eye’s visual field taken as 163° (Easter Jr & Nicola, 1996).

In Figure 4, linear fits between pYaw, pTrans and pConv components and pre-saccadic prey position were made using the MATLAB fitlm function with robust fit option. Prey position was calculated in gaze-referenced space at convergent saccade onset. To do this, the angle between the vector connecting the midpoint of the eyes and the prey target and the orientation of the head was computed and then the angle of the most proximal point of binocular overlap was subtracted.

### Latency and velocity main sequence analyses

Saccade initiation time, velocity and amplitude estimates were refined prior to analysis of inter-eye and eye-tail latency (Figure 5) and velocity main sequence (Figure 6). To estimate initiation times, we first computed two eye velocity estimates (Ev and Evsmooth) using raw eye position data up-sampled onto a 500 Hz time-base. Ev was determined by first smoothing eye position with the same custom LOWESS function described above and then computing the time derivative using the MATLAB gradient function. Evsmooth was determined by smoothing eye position with the same custom LOWESS function followed by an additional LOWESS function of span 50 ms, followed by computing the time derivative. The saccade midpoint was defined as the time-point at which Evsmooth peaked within a 160 ms window beginning 20 ms prior to the initial estimate of saccade onset time (see above). Then, saccade initiation was defined as the first time-point where Ev exceeded 20°/s prior to the saccade midpoint.

Saccade amplitude was calculated as the difference between pre- and post-saccadic eye position. Pre-saccadic eye position was the eye position one time-point (2 ms) prior to saccade initiation. Post-saccadic eye position was the median eye position over a 200 ms window starting at the time-point of peak eye displacement post-saccade. Peak displacement was the greatest change in eye position during a 200 ms window starting at saccade initiation, with the direction (positive or negative) determined by saccade type.

Peak eye velocity was maximum/minimum value of Ev during a time envelope that spanned the midpoint of the saccade. The envelope was defined by time-points where Evsmooth exceeded 10°/s.

For biphasic convergent saccades, the reversing eye was analysed as follows. Initiation of temporal eye movement was defined as the time-point at which Ev exceeded 20°/s in the temporal direction. Initiation of the second nasal rotation was defined as the time-point, following temporal movement initiation, at which Ev was closest to zero. Temporal velocity was the maximum Ev in the temporal direction between temporal and nasal initiation times and the amplitude of the temporal component was difference between eye position at these times. Velocity for the second nasal component was maximum Ev in the nasal direction during a 160 ms window following the initiation time of that movement; amplitude was the difference between post-saccadic eye position (as defined above) and eye position at second nasal movement onset.

Velocity main sequence relationships were fit using the function,

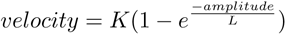

following Baloh et al. (1975), where *K* and *L* are constants. Fits were made using the MATLAB fitnlm function. We calculated the Aikake Information Criterion (AIC) to compare models with separate fits to each saccade type versus a single fit to data pooled across types. Fits were made with equal sample sizes from each saccade type (by randomly sampling from the type with more samples) and AIC values were normalised by dividing by the AIC value of the model with one exponential fit to the pooled data.

Linear velocity main sequence fits were made using the MATLAB fitlm function with robust options on and no bias term. For conjugate saccades, we only included saccades of amplitude *<* 10°, to limit the model to the non-saturating portion of the main sequence.

### Statistical analyses

All statistical analyses were performed in MATLAB. Types of statistical test and *N* are reported in the text or figure legends. All tests were two-tailed and we report p-values without correction for multiple comparisons unless otherwise noted.

---

**Figure S1:**
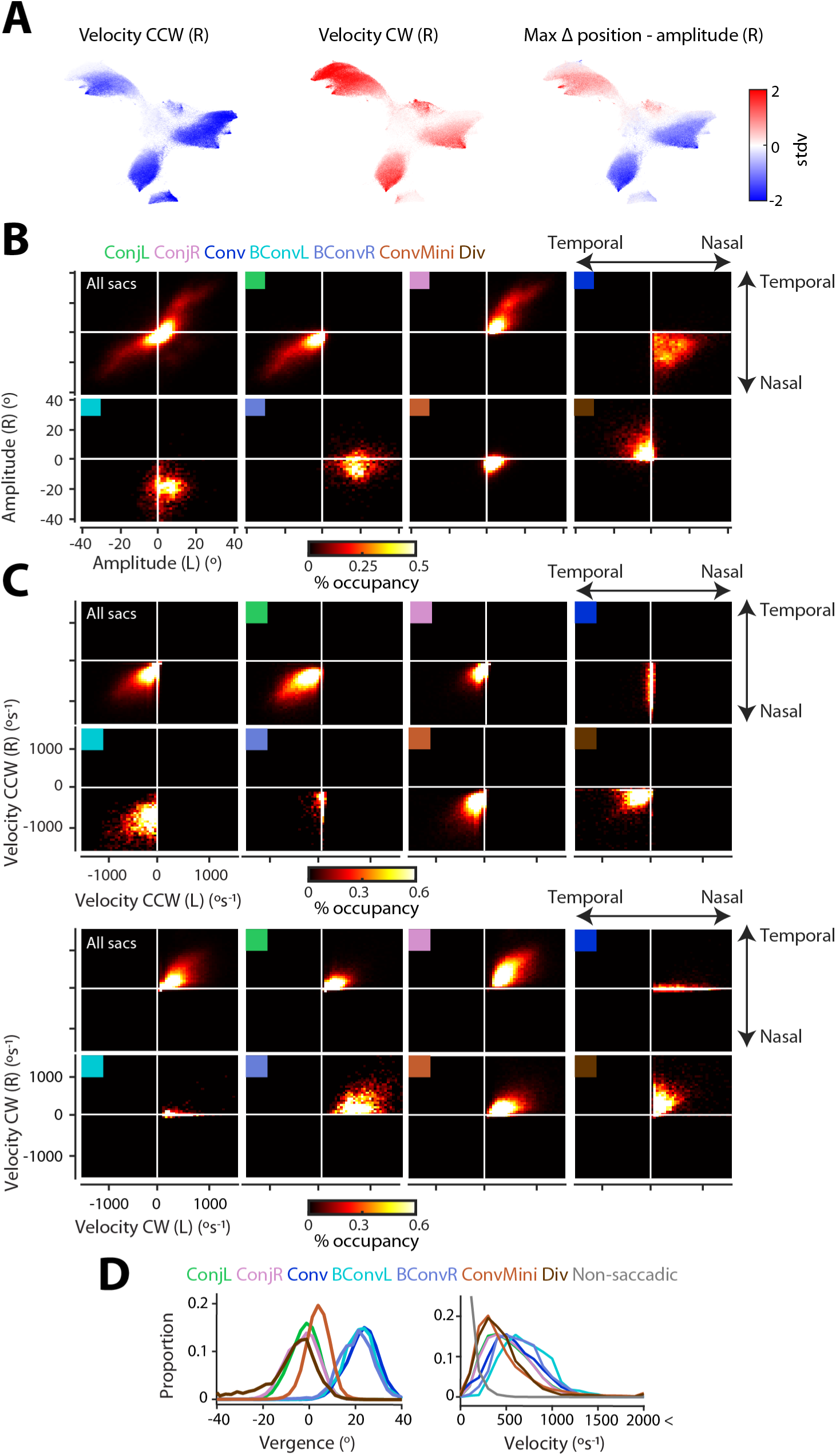
Additional saccade metrics from tethered larvae. Related to Figure 1. (A) UMAP embedding coloured according to three additional oculomotor metrics. (B–C) 2D histograms of saccade amplitude (B) and velocity (C). Saccade type indicated by coloured key in top left of each panel. (D) Post-saccadic vergence (left) and absolute eye velocity (right), across saccade types. For velocity histogram, non-saccadic cluster is included (grey, see Methods).

**Figure S2:**
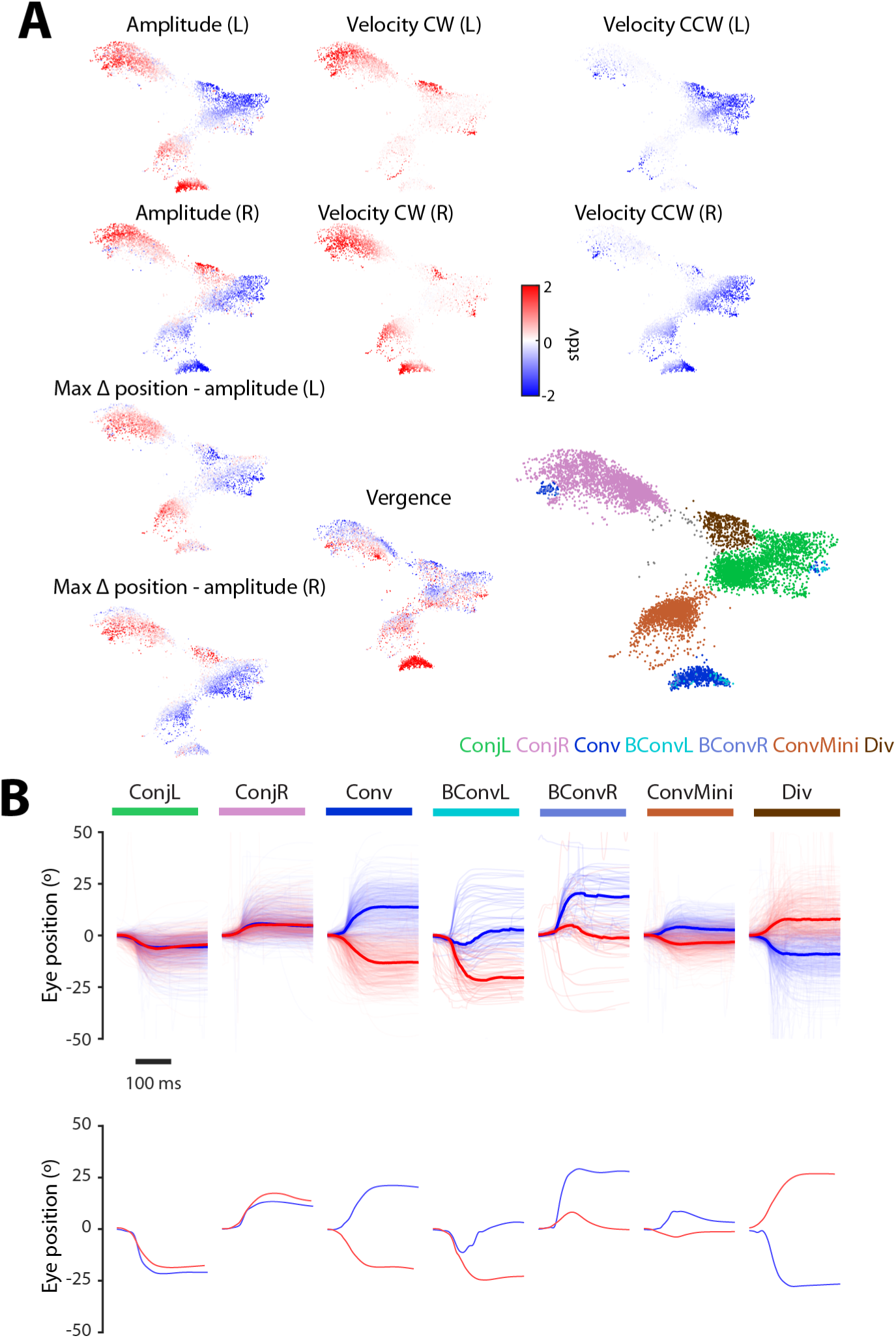
Saccade metrics from freely swimming fish. Related to Figure 1. (A) Rapid eye movements (9,367 events from 8 fish) after supervised embedding into the 2D UMAP space from Figure 1, coloured by normalised kinematic metrics and saccade type label. (B) *Top:* For each saccade type, 500 eye position traces are plotted with the median overlaid in bold. *Bottom:* A single example saccade from each type.

**Figure S3:**
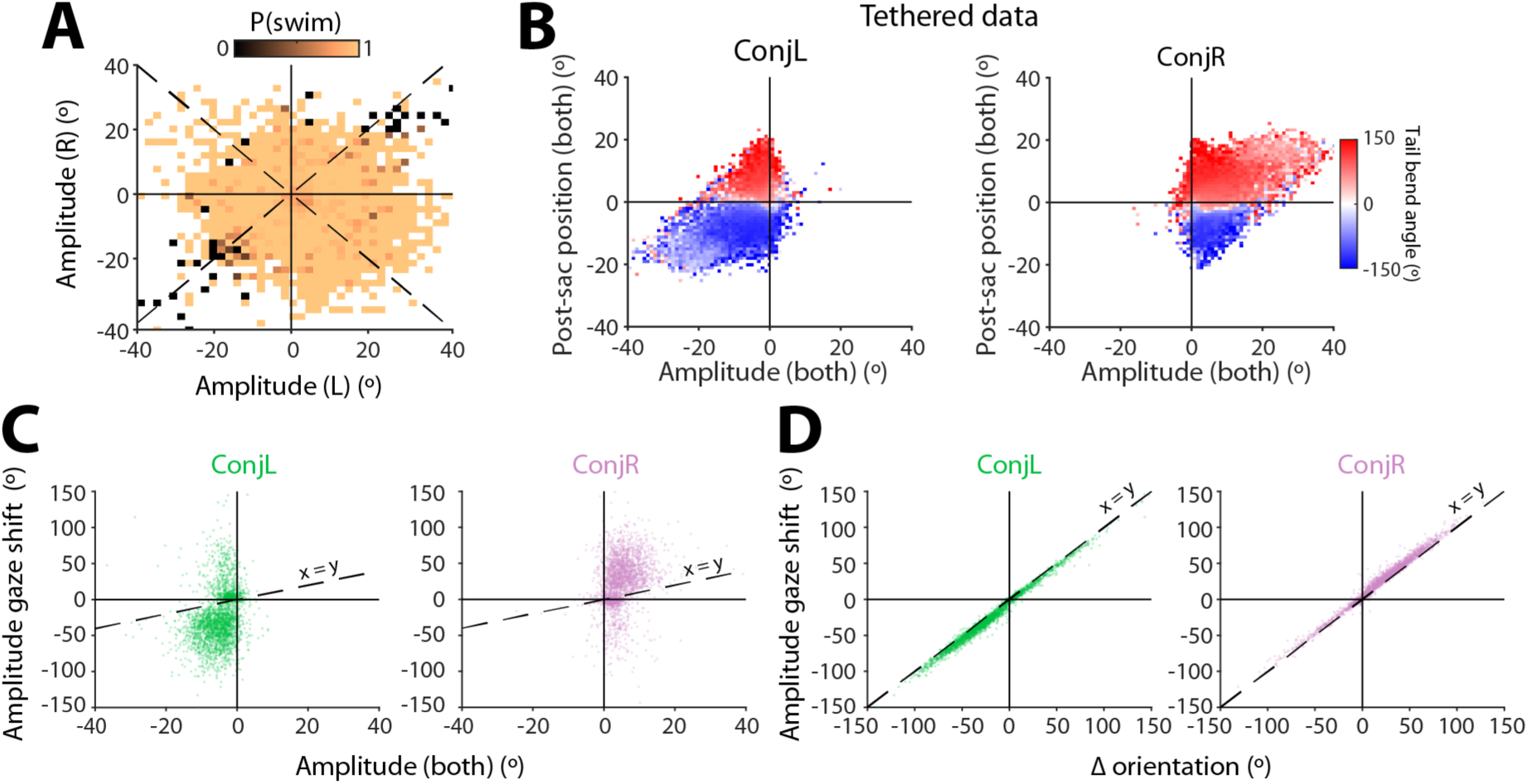
Conjugate saccades – additional data. Related to Figure 3. (A) Saccade amplitude coded by the probability of a coincident swim (data from 8 freely swimming larvae). (B) Left and right conjugate saccades binned by amplitude and post-saccadic conjugate eye position and colour-coded by median tail bend angle (152 tethered fish). (C–D) Amplitude of gaze shift versus change in conjugate eye position (C) or change in body orientation (D) (5,869 saccades from 8 freely swimming fish).

**Figure S4:**
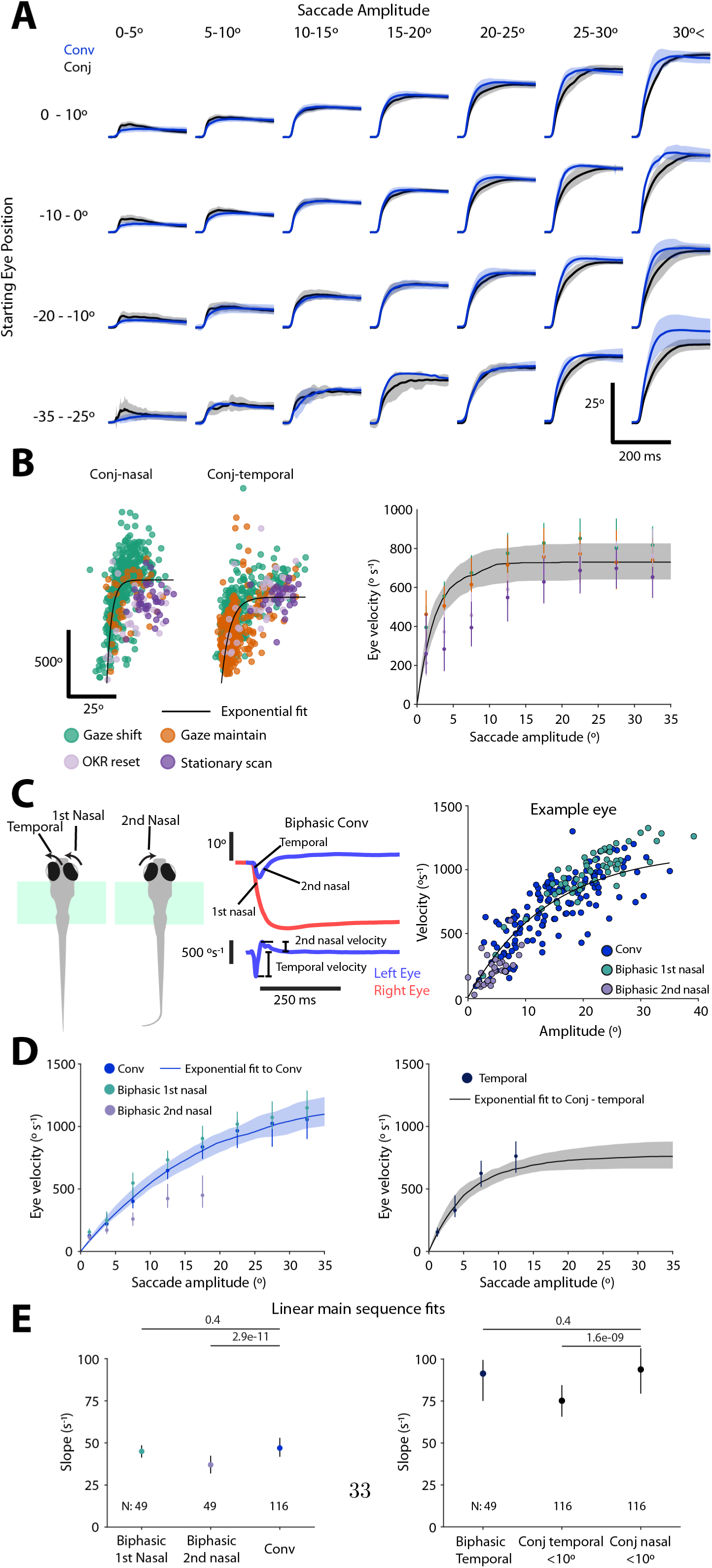
Velocity main sequence relationships – additional data. Related to Figure 6. (A) Eye position time series for adducting (nasal) saccadic eye movements from convergent and conjugate saccades of different amplitudes, binned by pre-saccadic eye position. (B) Velocity main sequence for sub-types of conjugate saccade. *Left:* An example eye with exponential fit and colour-coded sub-types of conjugate saccade. *Right:* Average velocity main sequence fit for conjugate saccades (reproduced from Figure 6D), overlaid with velocity data for each subtype of conjugate saccade (median ± IQR per amplitude bin). (C) *Left:* Illustration of biphasic convergent saccade, with component eye movements indicated. *Middle:* Position and velocity time series from an example biphasic convergent saccade. *Right:* Example eye showing exponential fit to regular convergent saccade data as well as components of biphasic saccades. (D) *Left:* Average velocity main sequence for regular convergent saccades (reproduced from Figure 6D), overlaid with velocity data for biphasic convergent saccades (median ± IQR per amplitude bin). *Right:* Average velocity main sequence for temporal eye movements within conjugate saccades, overlaid with velocity data for temporal components of biphasic convergent saccades (median ± IQR per amplitude bin). (E) Linear velocity main sequence fits. *Left:* Convergent saccades and nasal components of biphasic saccades. *Right:* Small amplitude (≤10°) conjugate saccades and temporal component of biphasic saccades. Linear fits made for eyes with at least 10 saccades (*N* eyes indicated). Fit slope coefficients plotted as median ± IQR. *p*-values from Kruskal-Wallis with Dunn-Sidak post-hoc tests.

## References

1. Antinucci, P., Folgueira, M., & Bianco, I. H. (2019). Pretectal neurons control hunting behaviour. Elife, 8.

2. Bahill, A. T., Bahill, K. A., Clark, M. R., & Stark, L. (1975a). Closely spaced saccades. Invest Ophthalmol, 14, 317–21.

3. Bahill, A. T., Clark, M. R., & Stark, L. (1975b). Computer simulation of overshoot in saccadic eye movements. Comput Programs Biomed, 4, 230–6.

4. Bahill, A. T., Clark, M. R., & Stark, L. (1975c). The main sequence, a tool for studying human eye movements. MATHEMATICAL BIOSCIENCES, 24, 191–204.

5. Bahill, A. T. & Troost, B. T. (1979). Types of saccadic eye movements. Neurology, 29, 1150–2.

6. Baloh, R. W., Sills, A. W., Kumley, W. E., & Honrubia, V. (1975). Quantitative measurement of saccade amplitude, duration, and velocity. Neurology, 25, 1065–70.

7. Bianco, I. H. & Engert, F. (2015). Visuomotor transformations underlying hunting behavior in zebrafish. Curr Biol, 25, 831–46.

8. Bianco, I. H., Kampff, A. R., & Engert, F. (2011). Prey capture behavior evoked by simple visual stimuli in larval zebrafish. Front Syst Neurosci, 5, 101.

9. Bianco, I. H., Ma, L.-H., Schoppik, D., Robson, D. N., Orger, M. B., Beck, J. C., Li, J. M., Schier, A. F., Engert, F., & Baker, R. (2012). The tangential nucleus controls a gravito-inertial vestibulo-ocular reflex. Curr Biol, 22, 1285–95.

10. Böhm, U. L., Prendergast, A., Djenoune, L., Nunes Figueiredo, S., Gomez, J., Stokes, C., Kaiser, S., Suster, M., Kawakami, K., Charpentier, M., Concordet, J.-P., Rio, J.-P., Del Bene, F., & Wyart, C. (2016). Csf-contacting neurons regulate locomotion by relaying mechanical stimuli to spinal circuits. Nat Commun, 7, 10866.

11. Bolton, A. D., Haesemeyer, M., Jordi, J., Schaechtle, U., Saad, F. A., Mansinghka, V. K., Tenenbaum, J. B., & Engert, F. (2019). Elements of a stochastic 3d prediction engine in larval zebrafish prey capture. Elife, 8.

12. Brainard, D. H. (1997). The psychophysics toolbox. Spat Vis, 10, 433–6.

13. Burgess, H. A. & Granato, M. (2007). Modulation of locomotor activity in larval zebrafish during light adaptation. J Exp Biol, 210, 2526–39.

14. Chen, C.-C., Bockisch, C. J., Straumann, D., & Huang, M. Y.-Y. (2016). Saccadic and postsaccadic disconjugacy in zebrafish larvae suggests independent eye movement control. Front Syst Neurosci, 10, 80.

15. Coubard, O. A. (2013). Saccade and vergence eye movements: a review of motor and premotor commands. Eur J Neurosci, 38, 3384–97.

16. Cullen, K. E. & Van Horn, M. R. (2011). The neural control of fast vs. slow vergence eye movements. Eur J Neurosci, 33, 2147–54.

17. Dunn, T. W., Gebhardt, C., Naumann, E. A., Riegler, C., Ahrens, M. B., Engert, F., & Del Bene, F. (2016a). Neural circuits underlying visually evoked escapes in larval zebrafish. Neuron, 89, 613–28.

18. Dunn, T. W., Mu, Y., Narayan, S., Randlett, O., Naumann, E. A., Yang, C.-T., Schier, A. F., Freeman, J., Engert, F., & Ahrens, M. B. (2016b). Brain-wide mapping of neural activity controlling zebrafish exploratory locomotion. Elife, 5, e12741.

19. Easter, S. S. & Johns, P. R. (1974). Horizontal compensatory eye movements in goldfish (carassius auratus). Journal of comparative physiology, 92, 37–57.

20. Easter, Jr, S. S. (1971). Spontaneous eye movements in restrained goldfish. Vision Res, 11, 333–42.

21. Easter Jr, S. S. & Nicola, G. N. (1996). The development of vision in the zebrafish (danio rerio). Developmental biology, 180, 646–663.

22. Ehrlich, D. E. & Schoppik, D. (2019). A primal role for the vestibular sense in the development of coordinated locomotion. Elife, 8.

23. Enright, J. T. (1984). Changes in vergence mediated by saccades. J Physiol, 350, 9–31.

24. Ester, M., Kriegel, H., Sander, J., & Xu, X. (1996). A density-based algorithm for discovering clusters in large spatial databases with noise. In Proceedings of the Second International Conference on Knowledge Discovery and Data Mining (KDD-96), Portland, Oregon, USA, E. Simoudis, J. Han, & U. M. Fayyad, eds., pp. 226–231. (AAAI Press).

25. Evinger, C. & Baker, R. (1991). Are there subdivisions of extraocular motoneuronal pools that can be controlled separately? Motor Control: Concepts and Issues.

26. Freedman, E. G. (2008). Coordination of the eyes and head during visual orienting. Exp Brain Res, 190, 369–87.

27. Freeman, J., Vladimirov, N., Kawashima, T., Mu, Y., Sofroniew, N. J., Bennett, D. V., Rosen, J., Yang, C.-T., Looger, L. L., & Ahrens, M. B. (2014). Mapping brain activity at scale with cluster computing. Nat Methods, 11, 941–50.

28. Friedrich, R. W., Jacobson, G. A., & Zhu, P. (2010). Circuit neuroscience in zebrafish. Curr Biol, 20, R371–81.

29. Gibaldi, A. & Sabatini, S. P. (2021). The saccade main sequence revised: A fast and repeatable tool for oculomotor analysis. Behav Res Methods, 53, 167–187.

30. Gremmler, S. & Lappe, M. (2017). Saccadic suppression during voluntary versus reactive saccades. J Vis, 17, 8.

31. Harris, A. J. (1965). Eye movements of the dogfish squalus acanthias l. J Exp Biol, 43, 107–38.

32. Henriques, P. M., Rahman, N., Jackson, S. E., & Bianco, I. H. (2019). Nucleus isthmi is required to sustain target pursuit during visually guided prey-catching. Curr Biol, 29, 1771–1786.e5.

33. Hermann, H. T. & Constantine, M. M. (1971). Eye movements in the goldfish. Vision Res, 11, 313–31.

34. Hernández, R., Calvo, P. M., Blumer, R., de la Cruz, R. R., & Pastor, A. M. (2019). Functional diversity of motoneurons in the oculomotor system. Proc Natl Acad Sci U S A, 116, 3837–3846.

35. Higashijima, S., Hotta, Y., & Okamoto, H. (2000). Visualization of cranial motor neurons in live transgenic zebrafish expressing green fluorescent protein under the control of the islet-1 promoter/enhancer. J Neurosci, 20, 206–18.

36. Horn, A. K. E. & Straka, H. (2021). Functional organization of extraocular motoneurons and eye muscles. Annu Rev Vis Sci, 7, 793–825.

37. Huang, Y.-Y. & Neuhauss, S. C. F. (2008). The optokinetic response in zebrafish and its applications. Front Biosci, 13, 1899–916.

38. Johnson, R. E., Linderman, S., Panier, T., Wee, C. L., Song, E., Herrera, K. J., Miller, A., & Engert, F. (2020). Probabilistic models of larval zebrafish behavior reveal structure on many scales. Curr Biol, 30, 70–82.e4.

39. Kapoula, Z. A., Robinson, D. A., & Hain, T. C. (1986). Motion of the eye immediately after a saccade. Exp Brain Res, 61, 386–94.

40. Kimura, Y., Satou, C., Fujioka, S., Shoji, W., Umeda, K., Ishizuka, T., Yawo, H., & Higashijima, S.-i. (2013). Hindbrain v2a neurons in the excitation of spinal locomotor circuits during zebrafish swimming. Curr Biol, 23, 843–9.

41. King, W. M. (2011). Binocular coordination of eye movements–hering’s law of equal innervation or uniocular control? Eur J Neurosci, 33, 2139–46.

42. King, W. M. & Zhou, W. (2000). New ideas about binocular coordination of eye movements: is there a chameleon in the primate family tree? Anat Rec, 261, 153–61.

43. Land, M. (2019). Eye movements in man and other animals. Vision Res, 162, 1–7.

44. Leyden, C., Brysch, C., & Arrenberg, A. B. (2021). A distributed saccade-associated network encodes high velocity conjugate and monocular eye movements in the zebrafish hindbrain. Sci Rep, 11, 12644.

45. Lister, J. A., Robertson, C. P., Lepage, T., Johnson, S. L., & Raible, D. W. (1999). nacre encodes a zebrafish microphthalmia-related protein that regulates neural-crest-derived pigment cell fate. Development, 126, 3757–67.

46. Marques, J. C., Lackner, S., Félix, R., & Orger, M. B. (2018). Structure of the zebrafish locomotor repertoire revealed with unsupervised behavioral clustering. Curr Biol, 28, 181–195.e5.

47. McInnes, L., Healy, J., & Melville, J. (2018). Umap: Uniform manifold approximation and projection for dimension reduction. arXiv, 10.48550/ARXIV.1802.03426.

48. Mearns, D. S., Donovan, J. C., Fernandes, A. M., Semmelhack, J. L., & Baier, H. (2020). Deconstructing hunting behavior reveals a tightly coupled stimulus-response loop. Curr Biol, 30, 54–69.e9.

49. Meehan, C., Ebrahimian, J., Moore, W., & Meehan, S. (2022). www.mathworks.com/matlabcentral/fileexchange/71902. MATLAB Central.

50. Meyer, A. F., O’Keefe, J., & Poort, J. (2020). Two distinct types of eye-head coupling in freely moving mice. Curr Biol, 30, 2116–2130.e6.

51. Michaiel, A. M., Abe, E. T., & Niell, C. M. (2020). Dynamics of gaze control during prey capture in freely moving mice. Elife, 9.

52. Miri, A., Bhasin, B. J., Aksay, E. R. F., Tank, D. W., & Goldman, M. S. (2022). Oculomotor plant and neural dynamics suggest gaze control requires integration on distributed timescales. J Physiol, 600, 3837–3863.

53. Naumann, E. A., Fitzgerald, J. E., Dunn, T. W., Rihel, J., Sompolinsky, H., & Engert, F. (2016). From whole-brain data to functional circuit models: The zebrafish optomotor response. Cell, 167, 947–960.e20.

54. Neuhauss, S. C., Biehlmaier, O., Seeliger, M. W., Das, T., Kohler, K., Harris, W. A., & Baier, H. (1999). Genetic disorders of vision revealed by a behavioral screen of 400 essential loci in zebrafish. J Neurosci, 19, 8603–15.

55. Orger, M. B., Smear, M. C., Anstis, S. M., & Baier, H. (2000). Perception of fourier and non-fourier motion by larval zebrafish. Nat Neurosci, 3, 1128–33.

56. Quinet, J., Schultz, K., May, P. J., & Gamlin, P. D. (2020). Neural control of rapid binocular eye movements: Saccade-vergence burst neurons. Proc Natl Acad Sci U S A, 117, 29123–29132.

57. Ramirez, A. D. & Aksay, E. R. F. (2021). Ramp-to-threshold dynamics in a hindbrain population controls the timing of spontaneous saccades. Nat Commun, 12, 4145.

58. Robinson, D. A. (2022a). The behavior of motoneurons. Prog Brain Res, 267, 15–42.

59. Robinson, D. A. (2022b). The function and phylogeny of eye movements. Prog Brain Res, 267, 1–14.

60. Robinson, D. A. (2022c). Neurophysiology, pathology and models of rapid eye movements. Prog Brain Res, 267, 287–317.

61. Robinson, D. A. (2022d). Properties of rapid eye movements. Prog Brain Res, 267, 271–286.

62. Salas, C., Herrero, L., Rodriguez, F., & Torres, B. (1997). Tectal codification of eye movements in goldfish studied by electrical microstimulation. f. Neuroscience, 78, 271–288.

63. Samonds, J. M., Geisler, W. S., & Priebe, N. J. (2018). Natural image and receptive field statistics predict saccade sizes. Nat Neurosci, 21, 1591–1599.

64. Schmitt, E. A. & Dowling, J. E. (1999). Early retinal development in the zebrafish, danio rerio: light and electron microscopic analyses. J Comp Neurol, 404, 515–36.

65. Schoonheim, P. J., Arrenberg, A. B., Del Bene, F., & Baier, H. (2010). Optogenetic localization and genetic perturbation of saccade-generating neurons in zebrafish. J Neurosci, 30, 7111–20.

66. Seideman, J. A., Stanford, T. R., & Salinas, E. (2018). Saccade metrics reflect decision-making dynamics during urgent choices. Nat Commun, 9, 2907.

67. Severi, K. E., Portugues, R., Marques, J. C., O’Malley, D. M., Orger, M. B., & Engert, F. (2014). Neural control and modulation of swimming speed in the larval zebrafish. Neuron.

68. Sharpe, J. A., Troost, B. T., Dell’Osso, L. F., & Daroff, R. B. (1975). Comparative velocities of different types of fast eye movements in man. Invest Ophthalmol, 14, 689–92.

69. Smit, A. C., Van Gisbergen, J. A., & Cools, A. R. (1987). A parametric analysis of human saccades in different experimental paradigms. Vision Res, 27, 1745–62.

70. Sparks, D. L. (2002). The brainstem control of saccadic eye movements. Nat Rev Neurosci, 3, 952–64.

71. Straka, H., Lambert, F. M., & Simmers, J. (2022). Role of locomotor efference copy in vertebrate gaze stabilization. Front Neural Circuits, 16, 1040070.

72. Sun, H. & Frost, B. J. (1998). Computation of different optical variables of looming objects in pigeon nucleus rotundus neurons. Nature neuroscience, 1, 296.

73. Sylvestre, P. A., Choi, J. T. L., & Cullen, K. E. (2003). Discharge dynamics of oculomotor neural integrator neurons during conjugate and disjunctive saccades and fixation. J Neurophysiol, 90, 739– 54.

74. Trivedi, C. A. & Bollmann, J. H. (2013). Visually driven chaining of elementary swim patterns into a goal-directed motor sequence: a virtual reality study of zebrafish prey capture. Front Neural Circuits, 7, 86.

75. Wolf, S., Dubreuil, A. M., Bertoni, T., Böhm, U. L., Bormuth, V., Candelier, R., Karpenko, S., Hildebrand, D. G. C., Bianco, I. H., Monasson, R., & Debrégeas, G. (2017). Sensorimotor computation underlying phototaxis in zebrafish. Nat Commun, 8, 651.

76. Yarbus, A. L. (1967). Eye Movements and Vision. (Plenum Press, New York).

77. Yoshimatsu, T., Schröder, C., Nevala, N. E., Berens, P., & Baden, T. (2020). Fovea-like photoreceptor specializations underlie single uv cone driven prey-capture behavior in zebrafish. Neuron, 107, 320– 337.e6.

78. Zhou, W. & King, W. M. (1998). Premotor commands encode monocular eye movements. Nature, 393, 692–5.

79. Zylbertal, A. & Bianco, I. H. (2023). Recurrent network interactions explain tectal response variability and experience-dependent behavior. Elife, 12.

